# Outer surface lipoproteins from the Lyme disease spirochete exploit the molecular switch mechanism of the complement protease C1s

**DOI:** 10.1101/2022.08.17.504303

**Authors:** Ryan J. Garrigues, Sheila Thomas, John M. Leong, Brandon L. Garcia

## Abstract

Proteolytic cascades comprise several important physiological systems, including a primary arm of innate immunity called the complement cascade. To safeguard against complement-mediated attack, the etiologic agent of Lyme disease, *Borreliella burgdorferi*, produces numerous outer surface-localized lipoproteins that contribute to successful complement evasion. Recently, we discovered a pair of *B. burgdorferi* surface lipoproteins of the OspEF-related protein family – termed ElpB and ElpQ – that inhibit antibody-mediated complement activation. In this study, we investigate the molecular mechanism of ElpB and ElpQ complement inhibition using an array of biochemical and biophysical approaches. *In vitro* assays of complement activation show that an independently folded homologous C-terminal domain of each Elp protein maintains full complement inhibitory activity and selectively inhibits the classical pathway. Using surface plasmon resonance, Alpha bead-based technology, and C1s enzyme assays, we show that binding of Elp proteins to activated C1s blocks C4 cleavage by competing with C1s/C4 binding without occluding the active site. C1s-mediated C4 cleavage is dependent on activation-induced binding sites, termed exosites. To test whether these exosites are involved in Elp/C1s binding, we performed site-directed mutagenesis which showed that ElpB– and ElpQ-binding require C1s residues in the anion-binding exosite located on the serine protease domain of C1s. Based on these results, we propose a model whereby ElpB and ElpQ exploit activation-induced conformational changes that are normally important for C1s-mediated C4 cleavage. Our study expands the known complement evasion mechanisms of microbial pathogens and reveals a novel molecular mechanism for selective C1s inhibition by Lyme disease spirochetes.

## Introduction

A first line of defense against invading pathogens is an evolutionarily ancient proteolytic cascade known as the complement system (1–4). Pattern recognition proteins of the complement system detect molecular signatures at the microbial surface and start a series of proteolytic reactions that convert inert complement components into bioactive protein fragments. A self-amplification loop involving complement component C3 quickly leads to opsonization of the activating surface, release of powerful chemoattractants known as anaphylatoxins (*i.e.*, C3a and C5a), and formation of a pore-like complex called the membrane attack complex (*i.e.*, C5b-9) that can directly kill susceptible microbes (1–4).

There are three primary mechanisms by which the complement cascade initiates, and these serve as the basis for three canonical pathways, known as the classical pathway (CP), lectin pathway (LP), and the alternative pathway (AP) (1–4). The CP is initiated by a multiprotein complex termed the first component of complement, C1, which is made up of a pattern recognition protein (C1q), and two homologous serine proteases (C1r and C1s), arranged as a heterotetramer (*i.e.*, C1qr_2_s_2_). The C1 complex primarily recognizes antibody-bound antigens leading to autoactivation of C1r, followed by C1r cleavage of C1s. Activated C1s then specifically cleaves complement component C2 and C4, promoting the formation of the central enzyme complexes of the complement cascade on the activating surface, known as C3 and C5 convertases. These complexes act to convert complement components C3 and C5 into active fragments, leading to the downstream effector functions of complement. The LP is initiated by recognition of foreign carbohydrate structures by mannan-binding lectin (MBL), ficolins, and certain collectins, which recruit MBL-associated proteases (MASPs), that like C1s, cleave C2 and C4. Thus, the CP and LP intersect at the CP/LP C3 convertase (*i.e.*, C4b2b) causing proteolytic activation of C3 and ultimately, of C5. In contrast, the AP is initiated spontaneously under a continuous low-level solution reaction known as tick-over that can lead to the direct formation of the AP C3 convertases (*i.e.*, C3bBb) on nearby unprotected surfaces. In addition to acting as the initiating enzyme for the AP, the AP C3 convertase establishes a self-amplification loop for all three pathways that results in large amounts of C3 activation, and deposition of the central opsonin of the cascade, C3b, at the activating surface.

Proteolytic cascades like the complement system, underly several important physiological systems including hemostasis and innate immune defense. Central to these cascades are the activities of a relatively small set of serine proteases that must act on a restricted set of substrates at the right time and in the right location. A common mechanism for control of proteolytic cascades is the production of an initial protease form that has very low catalytic activity, known interchangeably as a proenzyme or zymogen. Zymogen proteases must themselves be proteolytically cleaved in order to become fully active enzymes. This zymogen conversion event leads to conformational changes in the protease that allow for productive interactions with the appropriate substrate. These changes include formation of pockets on the protease, known as subsites, that bind directly to residues of a specific peptide sequence in a substrate scissile loop (5). In some cases, including the complement serine protease C1s, it has been shown that zymogen conversion results in conformational changes outside of the active site leading to formation of what are known as exosites (6–8). Exosites are critical for mediating contact with large protein substrates, such as C4 (205 kDa), in the case of C1s (6, 7, 9). Thus, in addition to subsite/scissile loop interactions, exosites can provide an additional layer of substrate specificity to proteases (10). However, the conformational changes associated with zymogen activation and exosite formation also provides a potential opportunity for highly selective intervention of proteolytic cascades by therapeutic agents – or less desirably – by microbial pathogens.

C1s consists of six sequentially arranged domains beginning with an N-terminal complement C1r/C1s, Uegf, Bmp1 (CUB) domain (CUB1), followed by an epidermal growth factor-like domain (EGF), a second CUB domain (CUB2), a complement control protein (CCP) domain (CCP1), a second CCP domain (CCP2), and the C-terminal catalytic serine protease (SP) domain. C4 binds to the C-terminal domains of C1s (*i.e.* CCP1-CCP2-SP) using three primary binding sites: i) C1s subsites that interact with the C4 scissile loop, ii) an exosite on the SP domain known as the anion-binding exosite (ABE) that is predicted to bind sulfotyrosine residues on C4, and iii) a second exosite at the hinge region between CCP1-CCP2 known as the CCP exosite (CCPE) that is predicted to bind to the C345C domain of C4 (6, 7, 9). Importantly, the mechanism of C1s-mediated C4 cleavage has been shown to rely on a “molecular switch” whereby zymogen activation of C1s results in conformational rearrangement leading to the formation of the C4-binding sites in the activated form of the protease (7). This model is further supported by a crystal structure of activated mannan-binding lectin serine protease 2 (MASP-2) in complex with C4, and C1s mutagenesis of four positively charged residues within the ABE (*i.e.*, K575, R576, R581, and K583) (6, 7, 9). Thus, C4 cleavage by C1s is controlled, in part, by exosite formation on C1r-cleaved forms of the C1s protease.

Activation of complement is proinflammatory, and is thus under tight control by proteins that are themselves part of the complement system, known as regulators of complement activity (RCAs) (11, 12). Microbial pathogens are also under pressure to evade complement recognition and a wide range of microbial complement evasion mechanisms have been described for medically important microbes including bacterial pathogens such as *Escherichia coli*, *Neisseria gonorrhoeae*, *Pseudomonas aeruginosa*, *Staphylococcus aureus*, *Streptococcus pneumoniae*, and the causative agent of Lyme disease, *Borreliella burgdorferi* (13, 14). Lyme disease is a tick-borne illness that is estimated to cause nearly half a million infections per year in the United States (15), many of which manifest in immunocompetent individuals. *B. burgdorferi*, has evolved a myriad of host immune evasion strategies, including a large anti-complement arsenal in the form of outer surface-localized lipoproteins (16, 17). Amazingly, over 10% of the surface lipoproteome of *B. burgdorferi* has been shown to interact directly with components of the complement system (16), and this percentage is even greater if one considers only the lipoproteins that are expressed during the vertebrate phase of infection. Thus, the Lyme disease spirochete has emerged as an important model to study evasion of the complement proteolytic cascade by microbial pathogens (16). While a majority of these *B. burgdorferi* lipoproteins function by recruiting the endogenous complement regulator factor H to the bacterial surface (18–26), or by targeting non-enzymatic components of the membrane attack complex (27), two separate classes of protease-targeting inhibitors have been identified in *B. burgdorferi*: i) BBK32 (28–30) and ii) ElpB/Q (31).

Using a surface lipoproteome library (32), we recently identified a new player in the *B. burgdorferi* complement evasion system in the form of two paralogous outer surface lipoproteins of the OspE/F-related protein family, termed ElpB and ElpQ (31) (also termed ErpB and ErpQ, respectively (33–35)). Genes encoding Elps are located on 32-kilobase circular plasmids (i.e. cp32 genetic elements) and are expressed during the mammalian phase of infection, suggesting a potential role in host interaction, which is further supported by the identification of laminin as a binding partner for ErpX/ElpX (35–39). We showed that ElpB and ElpQ bound with high affinity to the CP proteases C1r and C1s and were selective for the activated forms of each protease (31). Unlike BBK32, binding of activated C1r did not directly block its cleavage of zymogen C1s.

Instead, we found that ElpB and ElpQ block the CP by binding directly to C1s and inhibiting cleavage of its two natural substrates, C2 and C4 (31). Along with *B. burgdorferi* BBK32, which binds and inhibits both zymogen and active forms of C1r (28), ElpB and ElpQ may protect Lyme disease spirochetes from antibody-mediated complement killing during host infection. However, the molecular mechanism by which ElpB and ElpQ block the classical pathway remains poorly understood. In this study, we set out to elucidate a specific mechanism that explains how the Elp/C1s interaction leads to selective CP blockade.

## Results

### An alpha-helical domain at the carboxy-terminus of ElpB and ElpQ inhibits the classical pathway of human complement

We have previously shown that residues 19-378 of ElpB and residues 19-343 of ElpQ, which constitute the mature proteins lacking the secretion and lipidation sequences, prevent the deposition of C4b in a serum-based ELISA assay that utilizes IgM as a specific activator of the classical pathway (31). To determine if this inhibitory activity could be isolated to a specific region of the Elp proteins, we designed, produced, and purified a set of twelve truncation mutants of ElpB and ElpQ. Each of these mutants were tested for their ability to dose-dependently inhibit in the classical pathway ELISA assay. An initial set of truncation mutants divided ElpQ into N-terminal residues 19-216 and C-terminal residues 217-343, however, both proteins failed to inhibit complement (**Fig. 1A**, green circles and teal circles, respectively). Interestingly, truncations involving longer C-terminal regions (residues 103-343, 168-343, and 181-343) all retained full-length-like ElpQ inhibitory activities, as judged by half-maximal inhibitory concentration (IC_50_) values (**Fig. 1A, Table 1**). Similarly, ElpB C-terminal truncation constructs composed of residues 111-378, 138-378, and 182-378 exhibited IC_50_ values similar to that of full-length ElpB (**Fig. 1B, Table 1**). As was observed for ElpQ, shorter C-terminal ElpB constructs lost all activity (*i.e.*, residues 243-378 and 288-378). Constructs for ElpB (residues 138-288) and ElpQ (residues 168-328) that removed 90 and 15 residues from the C-termini of ElpB and ElpQ, respectively, produced inhibitors with weaker relative inhibitory activities (**Fig. 1A-B, Table 1**). This analysis defined a homologous C-terminal domain of ElpB and ElpQ that retains full complement inhibitory activities (*i.e.*, ElpB_182-378_ and ElpQ_181-343_) (**Fig. 1C, S1A-B**). Circular dichroism spectra were obtained for these inhibitory fragments as well as their full-length counterparts. These experiments indicate that each protein is folded and has strong alpha helical character (**Fig. S2A-D**). As further evidence that these constructs retained full inhibitory activity, ElpQ_181-343_ and ElpB_182-378_ dose-dependently inhibited hemolysis of opsonized rabbit erythrocytes, indicating that they prevent the downstream classical pathway-mediated formation of the membrane attack complex as demonstrated previously for full-length proteins (31) (**Fig. S2E**). Importantly, the inhibitory activity of both full-length and truncated Elp proteins were selective for the CP (**Fig. 1D-E**). Collectively, these experiments demonstrate that an independently folding domain at the C-terminus of ElpB and ElpQ selectively inhibits the CP in a manner similar to that of the full-length proteins.

**Figure 1.**
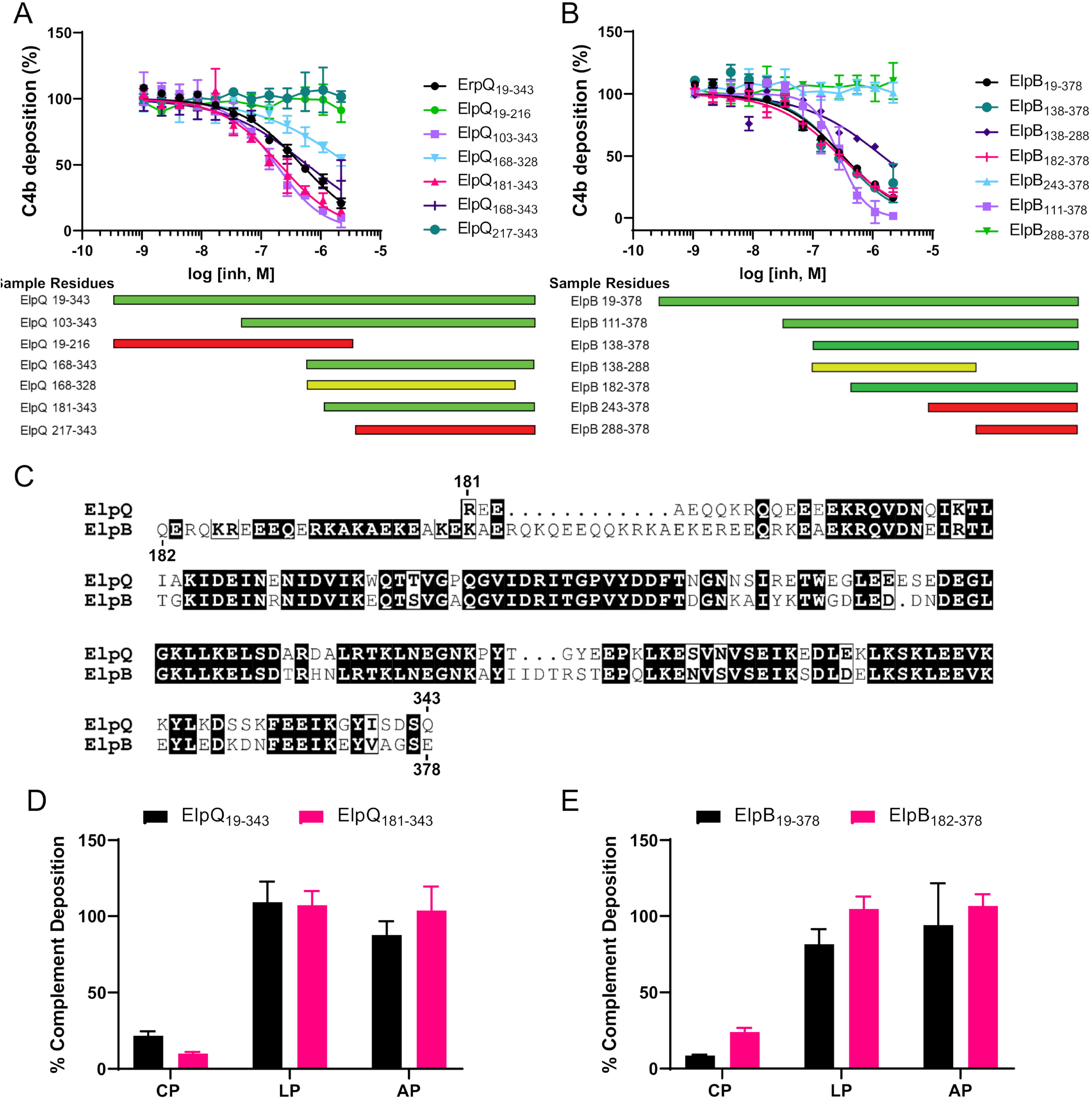
Complement inhibition assays using truncations of ElpQ and ElpB. **A-B**) Classical pathway ELISA inhibition assays were performed using C4b deposition as a marker for complement activation. A concentration series of each ElpQ and ElpB truncation protein were evaluated in triplicate. IC_50_ values are presented in **Table 1**. A qualitative schematic of ElpQ and ElpB truncation inhibitory activities are shown on the bottom panels: wild-type-like inhibition (green), non-wild-type-like inhibition (yellow), and no detectable inhibition (red). **C)** Sequence alignment of ElpQ_181-343_ and ElpB_182-378_ with the N-terminal and C-terminal residues numbers indicated above (ElpQ) and below (ElpB). Alignment was performed with EMBOSS Needle and rendering performed with ESPript 3.0 (61, 62). **D-E)** 2 μM concentration of **D)** ElpQ or **E)** ElpB proteins in classical (CP), lectin (LP), and alternative pathway (AP) ELISA complement activation assays. Deposition products C4b (CP/LP) and C3b (AP) are presented as a function of no inhibitor controls.

**Table 1.** CP ELISA IC50 values and confidence intervals.

| **<u>ElpQ Proteins</u>** | **<u>IC<sub>50</sub> (nM)</u>** | **<u>95% CI (nM)</u>** | **<u>ElpB Proteins</u>** | **<u>IC<sub>50</sub> (nM)</u>** | **<u>95% CI (nM)</u>** |
| --- | --- | --- | --- | --- | --- |
| ElpQ<sub>19-343</sub> | 480 | 420 to 550 | ElpB<sub>19-378</sub> | 320 | 280 to 370 |
| ElpQ<sub>103-343</sub> | 180 | 150 to 220 | ElpB<sub>111-378</sub> | 280 | 230 to 330 |
| ElpQ<sub>217-343</sub> | n.d. | n.d. | ElpB<sub>243-378</sub> | n.d. | n.d. |
| ElpQ<sub>168-343</sub> | 590 | 460 to 780 | ElpB<sub>288-378</sub> | n.d. | n.d. |
| ElpQ<sub>181-343</sub> | 210 | 170 to 260 | ElpB<sub>138-288</sub> | 1500 | 1100 to 2300 |
| ElpQ<sub>168-328</sub> | 3000 | 2000 to 5900 | ElpB<sub>138-378</sub> | 300 | 220 to 430 |
| ElpQ<sub>19-216</sub> | n.d. | n.d. | ElpB<sub>182-378</sub> | 290 | 250 to 330 |

n.d. = not determined, CI = confidence interval, IC<sub>50</sub> = half-maximal inhibitory concentration

### ElpB_182-378_ and ElpQ_181-343_ bind human C1s with high affinity

We have previously correlated the complement inhibition properties of full-length ElpB and ElpQ to binding of activated forms of the protease C1s, which is consistent with the pathway specificity observed above (**Fig. 1D-E**) (31). To test if the minimal inhibitory fragments of ElpB and ElpQ retained high-affinity binding to activated C1s, we used surface plasmon resonance (SPR). Biosensors were generated by immobilizing ElpB_182-378_ and ElpQ_181-343_ on an SPR sensorchip and purified activated C1s enzyme was titrated over each surface. The resulting sensorgrams were used to calculate equilibrium dissociation constants (*K*_D_) by fitting the data to a steady-state (*K*_D, ss_) and a kinetic binding model (*K*_D, kin_) (**Table 2**). In these experiments ElpQ_181-343_ bound purified activated C1s with *K*_D, kin_ = 17 ± 0.5 nM (**Table 2**). Similarly, the fully inhibitory ElpB_182-378_ truncation bound purified activated C1s with a calculated *K*_D, kin_ = 47 ± 3.1 nM. The values are comparable to previously obtained values for C1s affinity for GST-tagged full-length ElpQ (4.5 nM) and ElpB (3.9 nM). As expected from their activity in the ELISA-based complement assay (**Fig. 1**), additional selected constructs that showed no inhibitory activity also failed to bind C1s (*i.e.*, ElpQ_19-216_ and ElpB_288-378_) (**Fig. S3A-B**), while the inhibitory construct ElpB_111-378_ retained high-affinity binding to activated C1s (*K*_D, kin_ = 21 ± 0.2 nM) (**Fig. S3C**). These data confirm that the C-terminal domains of ElpB and ElpQ retain binding activity for activated forms of human C1s and are consistent with the notion that C1s-binding is important for ElpB/Q-mediated complement inhibition.

**Table 2.** SPR parameters

| **<u>Ligand</u>** | **<u>Analyte</u>** | **<u>$K_{D, \text{kin}}$ (nM)</u>** | **<u>$K_{D, \text{ss}}$ (nM)</u>** |
| --- | --- | --- | --- |
| ElpQ<sub>181-343</sub> | C1s Enzyme | $17 \pm 0.5$ | $31 \pm 0.4$ |
| | C1s CCP1-CCP2-SP | n.d. | $210 \pm 0.5$ |
| | C1s CCP1-CCP2-SP CCPE | n.d. | $370 \pm 0.5$ |
|  | C1s CCP1-CCP2-SP ABE | n.d. | n.d. |
|  | C1s CCP1-CCP2-SP CCPE/ABE | n.d. | n.b. |
| ElpB<sub>182-378</sub> | C1s Enzyme | $47 \pm 3.1$ | $85 \pm 2.3$ |
| | C1s CCP1-CCP2-SP | n.d. | $360 \pm 17$ |
| | C1s CCP1-CCP2-SP CCPE | n.d. | $570 \pm 110$ |
|  | C1s CCP1-CCP2-SP ABE | n.d. | n.d. |
|  | C1s CCP1-CCP2-SP CCPE/ABE | n.d. | n.b. |
| ElpQ<sub>111-378</sub> | C1s Enzyme | $21 \pm 0.2$ | $35 \pm 0.4$ |

n.d. = not determined, n.b. = no binding detected, $K_{D, \text{kin}} = K_D$ calculated from kinetic fit, $K_{D, \text{ss}} = K_D$ calculated from steady-state fit (see *Experimental Procedures*)

### ElpB and ElpQ bind to the C-terminal region of C1s in a manner that depends on the activation state of the protease

We have previously shown that ElpB and ElpQ preferentially bind to the activated form of full-length C1s (31). Separately, we have shown that these inhibitors block cleavage of the large endogenous substrates of C1s (*i.e.*, C2 and C4), but not a small synthetic peptide substrate (31). In considering potential explanations for these results, we hypothesized that ElpB/Q may be targeting the C4-binding site on activated C1s located on the C-terminal CCP1-CCP2-SP domains (**Fig. 3A**). To begin to test this hypothesis, we produced several C1s truncations involving the C4-binding domains of C1s. Using full-length ElpQ as an initial model for ElpB/Q-binding, we evaluated the binding activity of these C1s derivatives by SPR (**Fig. 3B**). Single concentration injections (500 nM) of C1s truncations, lacking the SP domain (*i.e.*, CUB2-CCP1 and CUB2-CCP1-CCP2), failed to interact with the ElpQ biosensor (**Fig. 3B**, red and brown lines). C1r-activated C1s-CCP1-CCP2-SP bound directly to ElpQ (**Fig. 3B**, green line), although our later titration experiments indicated that the binding affinity (*K*_D, ss_ = 210 nM) was ~7-fold lower relative to that of full-length activated C1s (*K*_D, ss_ = 31 nM), suggesting that domains outside of CCP1-CCP2-SP contribute to Elp-binding (**Table 2**). Next, we found that ElpB also bound to CCP1-CCP2-SP. The binding affinity (*K*_D, ss_ = 360 nM) for the C-terminal fragment of C1s, like that of ElpQ, was somewhat lower (~4-fold) relative to that of full-length activated C1s (*K*_D, ss_ = 85 nM) (**Table 2**). Most strikingly, when the zymogen form of C1s-CCP1-CCP2-SP was injected, no interaction was detected (**Fig. 3B**, black line). This result is consistent with our previous observations that ElpQ preferentially bound activated full-length C1s (31). This result is also reminiscent of the observations made by Perry *et. al* where C4 failed to bind to zymogen C1s-CCP1-CCP2-SP, but bound strongly to activated C1s-CCP1-CCP2-SP, using a similar SPR-based approach (7).

**Figure 2.**
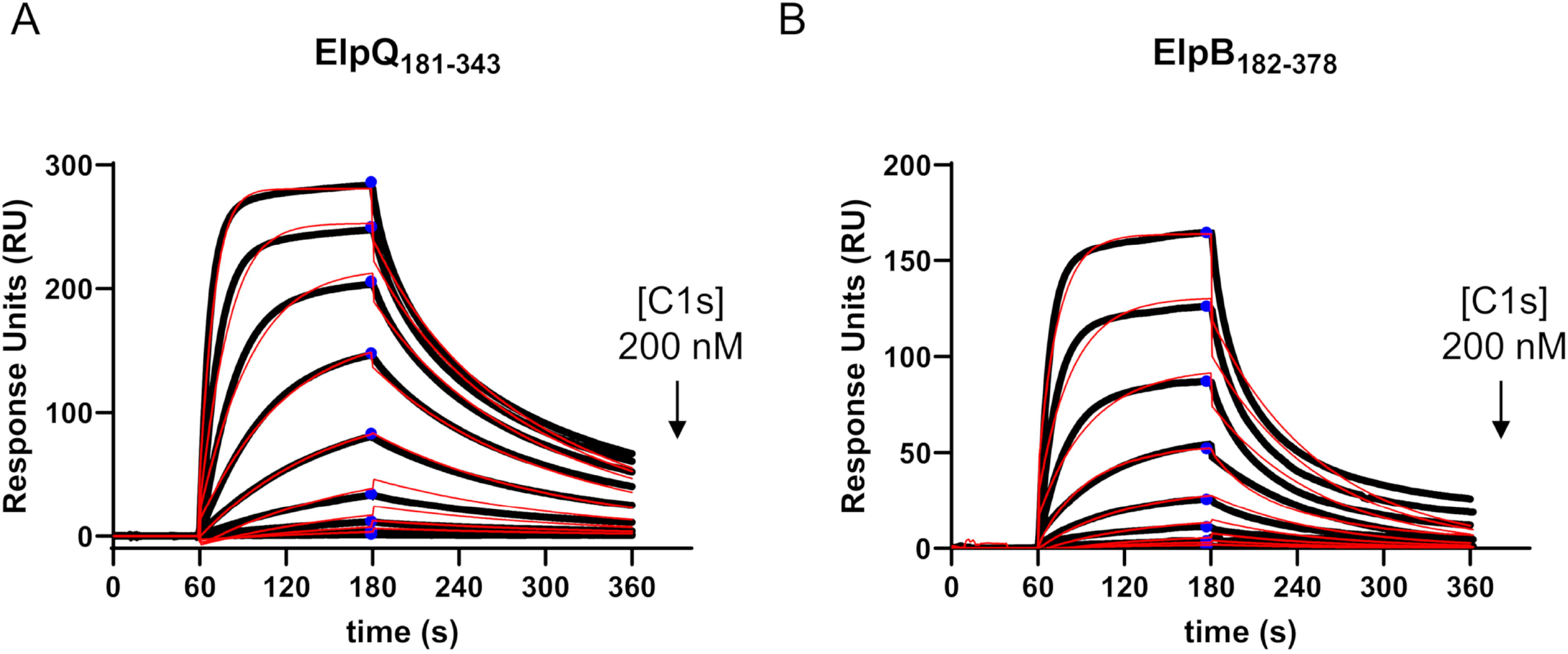
SPR binding assays using activated full-length C1s. **A-B)** ElpQ_181-343_ and ElpB_182-378_ were immobilized on an SPR sensor chip by amine coupling chemistry. Multi-cycle SPR experiments were performed using a two-fold dilution series (200 – 0.4 nM) of purified full-length activated C1s. Reference-subtracted sensorgrams are shown as black lines. A representative injection series is shown. Each experiment was performed in triplicate. Sensorgrams were fit to steady-state (*K*_D, ss_, blue circles) and kinetic models (*K*_D, kin_, red traces) of binding using Biacore T200 Evaluation software and calculated *K*_D_ values are shown in **Table 2**.

**Figure 3.**
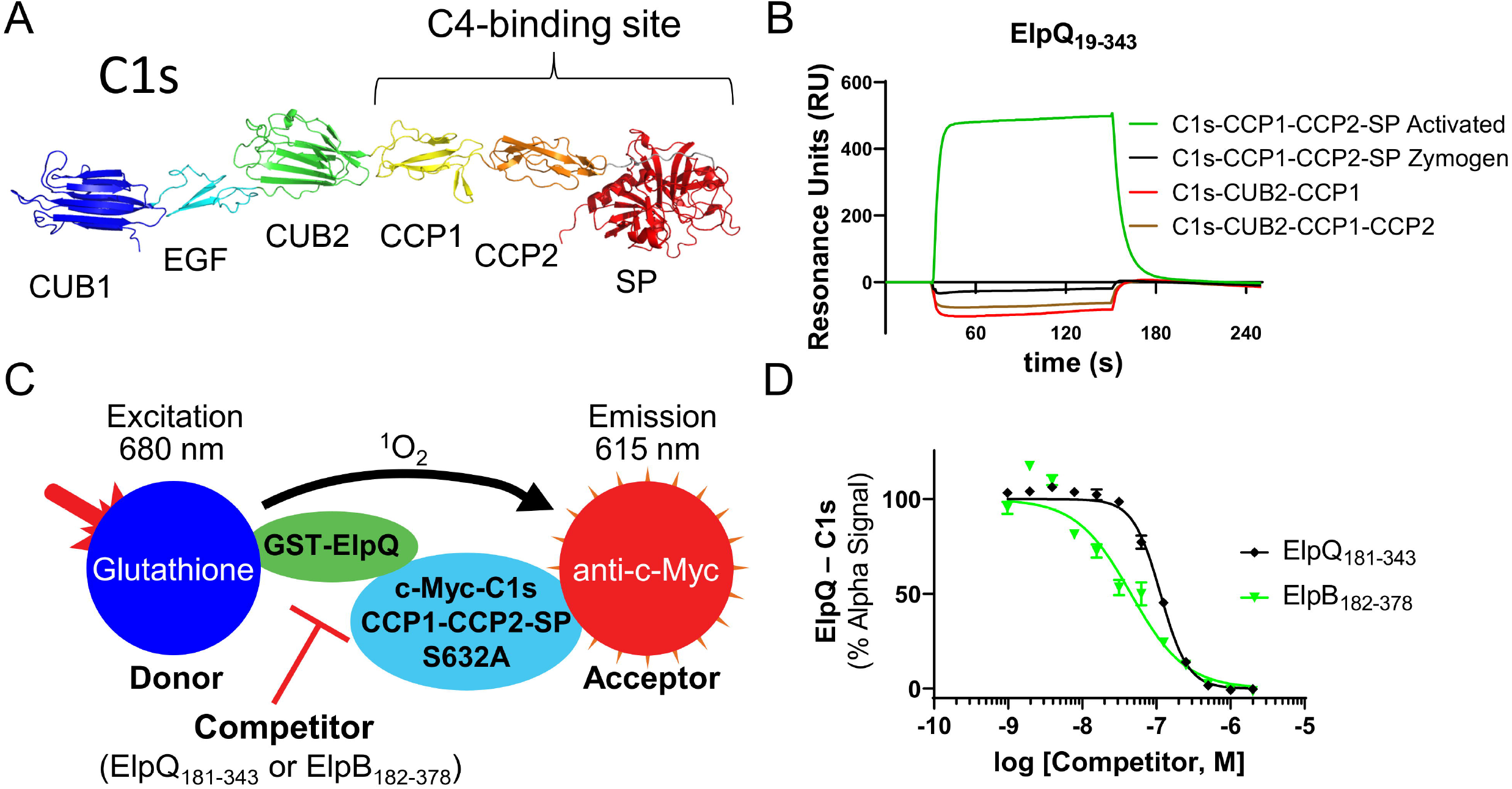
ElpQ and ElpB bind to the C-terminal domains of activated C1s. **A)** An AlphaFold (63) model of full-length human C1s lacking the N-terminal signal sequence is shown (UNIPROT #: P09871). The domain architecture of C1s and the C4-binding site on C1s are highlighted. **B)** ElpQ_19-343_ was immobilized on a SPR sensorchip. Selected C1s truncation proteins containing CUB2, CCP1, CCP2 and SP domains were injected over the ElpQ biosensor at 500 nM. Representative curves from a single injection are shown from an experiment performed in triplicate. **C)** Alpha competition experiment schematic. Donor beads conjugated with glutathione bound to 50 nM GST-ElpQ were added to anti-c-Myc antibody conjugated Acceptor beads bound to 500 nM activated c-Myc-C1s-CCP1-CCP2-SP_S632A_ produce an Alpha signal (**Fig. S4**). **D)** Untagged ElpQ_181-343_ or ElpB_182-378_ were titrated (2,200 – 1 nM) and incubated for 1 hr at room temperature in the dark. Alpha signal was obtained with excitation at 680 nm and emission was at 615 nm. Alpha signals were normalized to wells containing no competitor (100%) and no C1s (0%). Curves were fit using non-linear regression and IC_50_ values and 95% confidence intervals are reported in **Table 3**. Experiments were performed in triplicate.

To validate the interaction of the Elp truncation mutant proteins with activated C1s-CCP1-CCP2-SP and to more directly compare ElpB and ElpQ, we developed an Alpha-based competition experiment. Alpha is a luminescent bead-based assay whereby an amplified luminescent signal is produced when an Alpha Donor bead is brought in close proximity (<200 nm) to an Alpha Acceptor bead (40). We initially screened for an Alpha signal to detect binding between GST-tagged ElpQ_19-343_ and a c-Myc-tagged form of an activated, but catalytically inert, derivative of C1s (*i.e.*, C1r-cleaved, C1s-CCP1-CCP2-SP harboring a mutation (*i.e.*, S632A) that eliminates C1s enzymatic activity). Consistent with the SPR experiments (**Fig. 3B**), several conditions were identified that yielded an Alpha signal, confirming a direct interaction between GST-ElpQ_19-343_ and activated c-Myc-C1s-CCP1-CCP2-SP_S632A_ (**Fig. S4**).

**Table 3.** Alpha parameters

| **<u>Competitor</u>** | **<u>K<sub>D</sub> (nM)</u>** | **<u>95% CI (nM)</u>** |
| --- | --- | --- |
| ElpQ<sub>19-343</sub> | 190 | 170 to 200 |
| ElpQ<sub>181-343</sub> | 110 | 110 to 120 |
| ElpB<sub>182-378</sub> | 45 | 37 to 56 |
| C4 | 740 | 580 to 1000 |

CI = confidence interval

To assess the ability of ElpQ and ElpB truncation proteins to inhibit the ElpQ-C1s interaction in the above assay, we utilized an Alpha-based competition assay with optimized fixed concentrations of each protein component (**Fig. 3C**, green/cyan), anti-c-Myc Alpha Acceptor (**Fig. 3C**, red), and Glutathione Alpha Donor (**Fig. 3C**, blue) beads. By evaluating a concentration series of “cold” (*i.e.*, non-GST-tagged) ElpQ_181-343_, in this competition assay (**Fig. 3D**), we found that ElpQ_181-343_ dose-dependently competed with the c-Myc-C1s-CCP1-CCP2-SPS_632A_/GST-ElpQ_19-343_ interaction with a *K*_D_,_ElpQ(181-343)_ = 110 nM. Furthermore, a similar concentration series of “cold” ElpB_182-378_ truncation protein also blocked this interaction, with a *K*_D_,_ElpB(182-378)_ = 45 nM. Collectively, these results demonstrate: i) that ElpB and ElpQ have a binding site on activated C1s-CCP1-CCP2-SP (*i.e.*, the C4-binding region of C1s); ii) that the minimal inhibitory fragments identified here (*i.e.*, ElpQ_181-343_, and ElpB_182-378_) compete with full-length ElpQ for this site; and iii) suggest that ElpB and ElpQ have overlapping binding sites on C1s-CCP1-CCP2-SP.

### ElpQ_181-343_ blocks C1s-CCP1-CCP2-SP-mediated C4 cleavage and competes with C4 for C1s-CCP1-CCP2-SP binding

We have previously linked the direct binding of full-length ElpB and ElpQ to inhibition of C4 cleavage by purified full-length C1s, but not a synthetic peptide C1s substrate (31). We sought to confirm that this inhibitory profile is retained when C1s-CCP1-CCP2-SP was used as an enzyme, rather than full-length C1s. To test this, we used the minimal inhibitory domain of ElpQ (i.e., ElpQ_181-343_) as a model for ElpB/Q inhibitory activity, and performed an SDS-PAGE gel-based C4 cleavage assay using activated C1s-CCP1-CCP2-SP enzyme. Indeed, ElpQ_181-343_ was capable of dose-dependent inhibition of C4 cleavage by C1s-CCP1-CCP2 (**Fig. 4A-B**), with an IC_50_ = 3.6 μM, which is comparable to the 11 μM IC_50_ for inhibition of full-length C1s by full-length ElpQ (31). Furthermore, like in full-length ElpQ/full-length C1s experiments (31), ElpQ_181-343_ (25 μM) failed to inhibit cleavage of a small synthetic peptide by C1s-CCP1-CCP2-SP (**Fig. 4C**).

**Figure 4.**
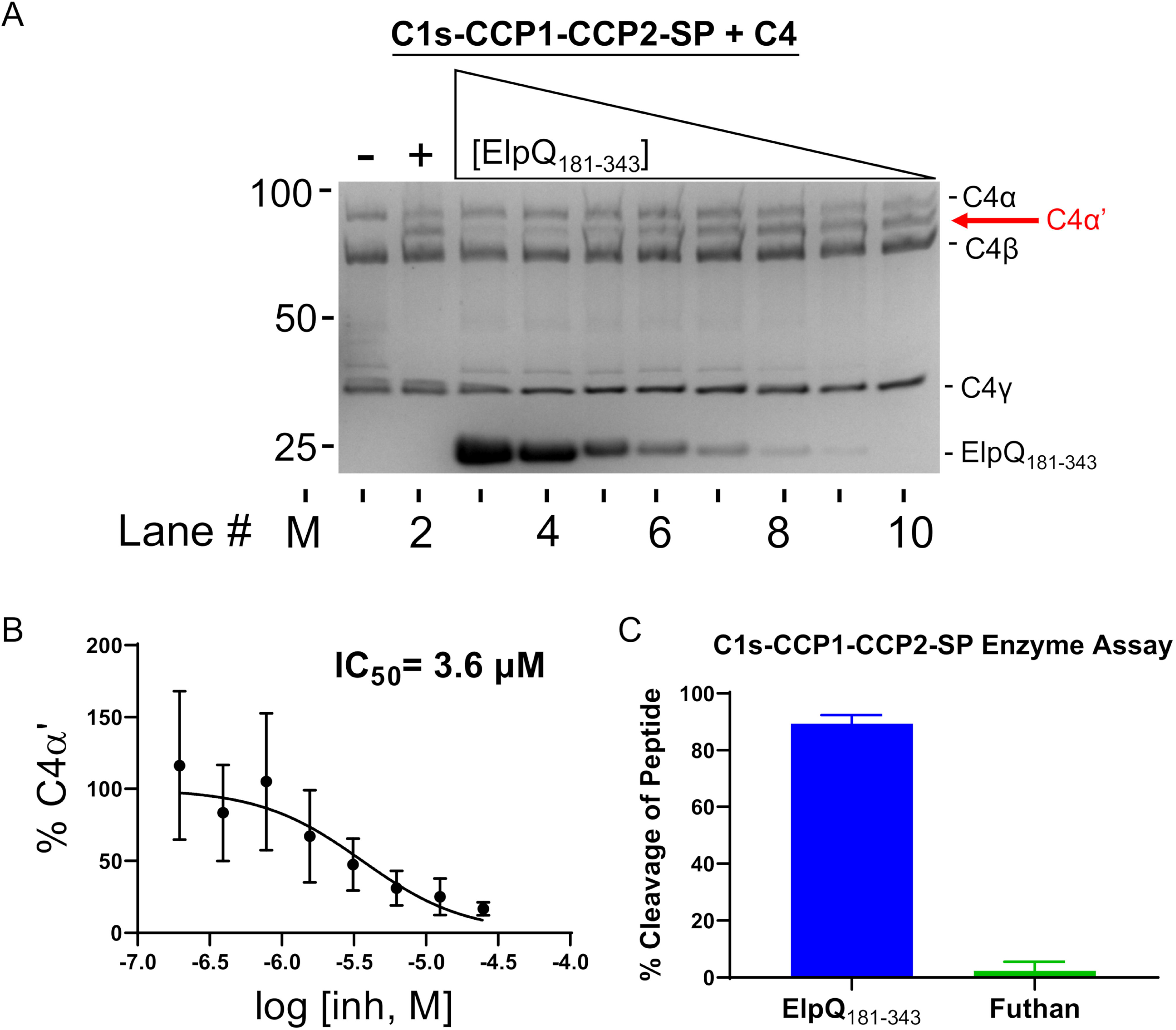
C1s-CCP1-CCP-SP enzyme inhibition assays. Enzymatic cleavage of C4 by C1s at 37° C for 1 hr results in four chains: C4α (~95 kDa), C4α’ (~85 kDa), C4β (~75 kDa), and C4γ (~35 kDa) that can be resolved by SDS-PAGE analysis. **A)** M: Marker (kDa). Lanes 1-2: SDS-PAGE gel of C4 cleavage in the presence (“+”) or absence (“−“) of 6.25 nM C1s-CCP1-CCP2-SP, and 620 nM C4. Lanes 3-10: C4 cleavage profile in the presence of 6.25 nM C1s, 620 nM C4, and a two-fold dilution series (from 25 to 0.20 μM) of ElpQ_181-343_. **B)** The fraction of C4α’ relative to input C4β in the same lane and normalized to C1s + C4 positive control (lane 2) and negative control C4 (lane 1) was determined by densitometry analysis. A representative gel of three independent experiments is shown. **C)** Enzymatic cleavage by C1s-CCP1-CCP2-SP of the small peptide substrate Z-L-Lys-sBzl was assayed with 5,5’-dithiobis-(2-nitrobenzoic acid) (Ellman’s reagent) in the presence of 25 μM ElpQ_181-343_ or Futhan (a small molecule C1s inhibitor). Absorbance was read at 412 nm and signals were normalized to negative control no-enzyme wells.

We hypothesized that Elp proteins block activated C1s-CCP1-CCP2-SP by interfering with the interaction of C4-binding to the C1s enzyme. To test this, we used our established Alpha-based competition assay and tested whether ElpQ competes with C4 for binding to activated C1s-CCP1-CCP2-SP_S632A_. Indeed, we observed that soluble C4 inhibited the interaction between GST-ElpQ_19-343_/activated c-Myc-C1s-CCP1-CCP2-SP_S632A_ in a dose-dependent manner with a *K*_D_ = 740 nM (**Fig. 5**). Untagged ElpQ_19-343_, used as a control, also competed, with a calculated *K*_D_ = 190 nM. This result demonstrates that ElpQ-binding of activated C1s-CCP1-CCP2-SP interferes with access of C4 to the C1s enzyme and provides mechanistic insight into the observed inhibitory effects in our gel-based C1s/C4 cleavage assay.

**Figure 5.**
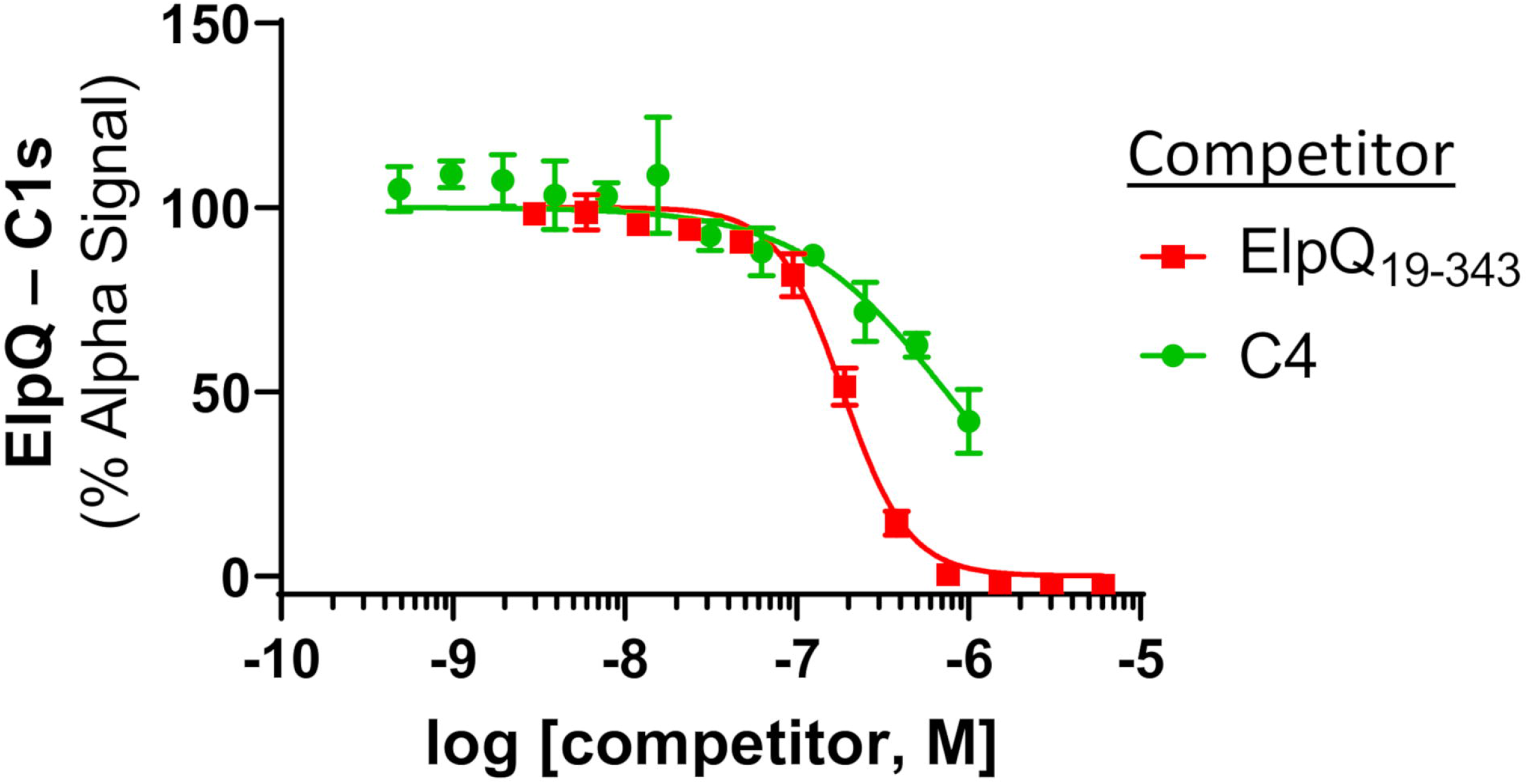
C4 competes with ElpQ binding of C1s-CCP1-CCP2-SP_S632A_. The Alpha competition experiment described in **Fig. 3C** was used to assess C4 competition. Purified C4 (1,000 – 0.5 nM) or an untagged ElpQ_19-343_ control (6,100 – 3 nM) were titrated and incubated for 1 hr at room temperature in the dark. Alpha signal was obtained with excitation at 680 nm and emission was at 615 nm. Alpha signals were normalized to wells containing no competitor (100%) and no C1s (0%). Curves were fit using non-linear regression. Experiments were performed in triplicate.

### High-affinity binding of ElpB and ElpQ to C1s-CCP1-CCP2-SP requires a known C4 exosite

It has been proposed that conversion of C1s from a zymogen form to an active form involves the formation of two distinct C4-binding exosites on activated C1s (**Fig. 6A, Fig. S5**). The data presented above suggests that ElpB and ElpQ may bind C1s in a C4-like manner and thus rely on interaction with one or both of the known C4 exosites on C1s. To test if C4 exosites on C1s are important for Elp interaction, we produced site-directed alanine mutants of the CCPE and ABE sites in activated C1s-CCP1-CCP2-SP. A construct was also created in which both sites were mutated (denoted “CCPE/ABE”). ElpQ_181-343_, and ElpB_182-378_ were tested for binding to these constructs using SPR. For each Elp derivative, mutation of CCPE alone resulted in a small (two-fold) difference in affinity, relative to wild-type C1s-CCP1-CCP2-SP (**Fig. 6B-D, Table 2**). In contrast, mutation of the ABE resulted in a construct of C1s with very weak binding such that *K*_D_ values could not be calculated (**Fig. 6B-D**). Furthermore, mutation of both exosites resulted in a complete loss of binding between C1s-CCP1-CCP2-SP and both Elp proteins (**Fig. 6B-D**). These data show that binding of C1s by ElpB or ElpQ requires C1s residues outside of the C1s active site that are known to be involved in C1s/C4 recognition. Taken together, our data suggest that ElpB and ElpQ bind activated C1s by targeting a primary exosite for C4 and thereby occlude its access and subsequent cleavage by the protease.

**Figure 6.**
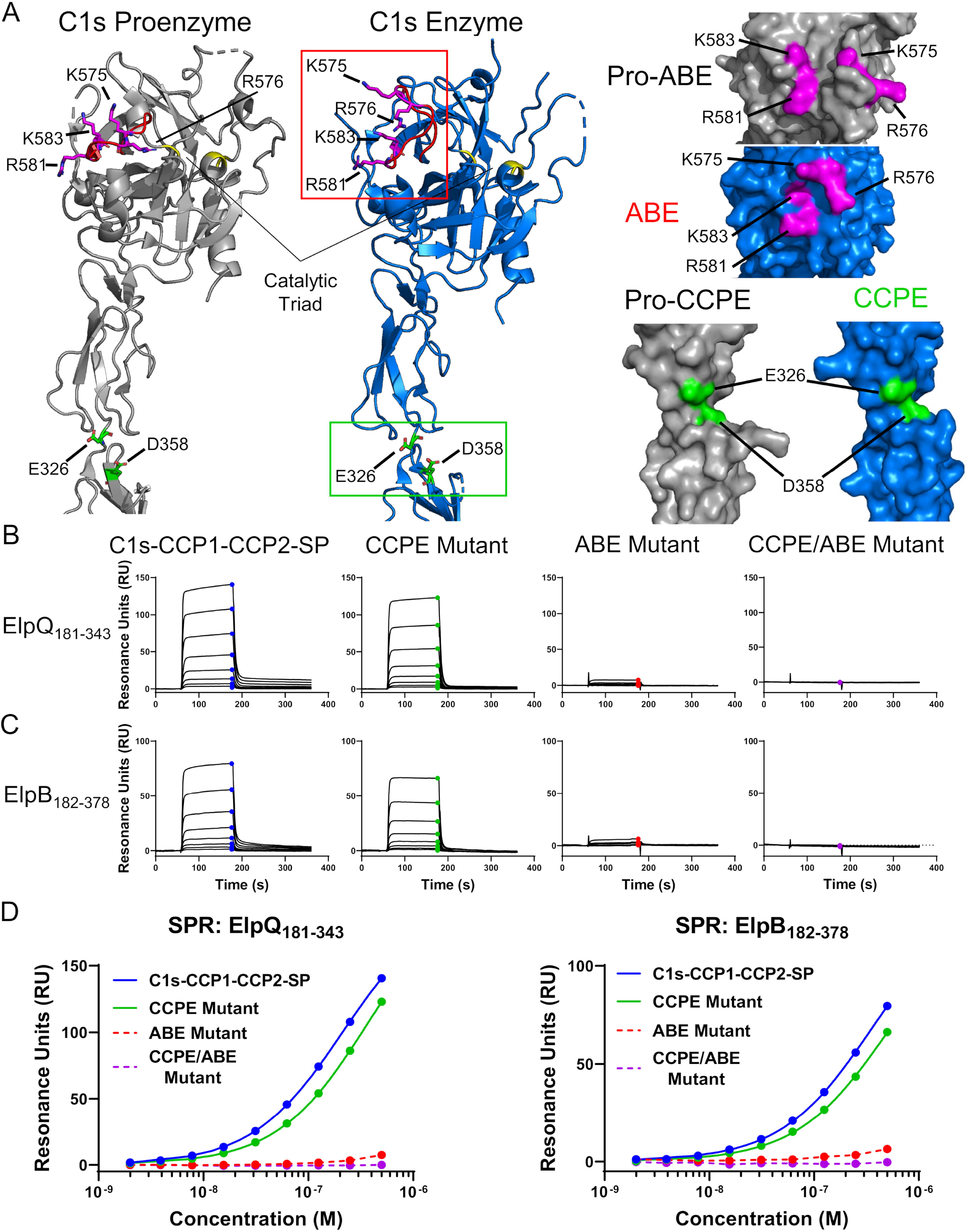
C1s-CCP1-CCP2-SP exosite mutagenesis SPR binding studies. **A)** Ribbon diagram of crystal structures of CCP1-CCP2-SP in a proenzyme (grey, PDB: 4J1Y) and activated form (blue, PDB: 5UBM). The catalytic triad is marked in yellow on each structure. Site-directed alanine mutagenesis was performed on residues in each of the C4 exosites: the anion binding exosite (ABE, red loop) consisting of K575-R576-R581-K583 (magenta sticks), the region between CCP1 and CCP2 termed the CCP exosite (CCPE) consisting of E326-D358 (green sticks), or simultaneous mutation of both (CCPE/ABE). Surface representations of each site, denoted proexosites for C1s proenzyme models, are shown from another vantage with the mutated residues highlighted. **B-C)** ElpQ_181-343_ and ElpB_182-378_ were immobilized on an SPR sensor chip and activated C1s-CCP1-CCP2-SP constructs were injected as a two-fold dilution curve (200 – 0.8 nM) symbols represent steady-state values at each concentration. **D)** Steady-state fits are shown of the raw sensorgrams presented in **B** and **C** with colored circles corresponding to the steady state RU. Representative curves are shown, and each injection series was performed in triplicate. Fits (solid lines) were performed using Biacore T200 Evaluation software and *K*_D_ values are presented in **Table 2**. ABE and CCPE/ABE mutants could not be fit and are shown as dashed lines.

## Discussion

Lyme disease spirochetes have evolved a powerful complement evasion system that includes the paralogous outer surface lipoproteins, ElpB and ElpQ (31). In the current study, we isolated the inhibitory activity of ElpB and ElpQ to a homologous C-terminal domain and showed that these truncations bind and inhibit C1s with similar overall activity to that of the full-length proteins (**Figs. 1, 2**). We note that the presence of a functional C-terminal domain is a common topology found in borrelial outer surface lipoproteins, including the C1r-binding domain of BBK32 (28, 41, 42). Like BBK32, the exosite-mediated inhibitory mechanism of ElpB and ElpQ described here is likely to provide the Lyme disease spirochete alternative means by which to thwart antibody-mediated complement killing (31). Interestingly, Lyme disease spirochetes produce several complement inhibitor proteins that have redundant targets but distinct inhibitory mechanisms (43). Target redundancy paired with mechanistic diversity is also seen in complement inhibitors from other notable bacterial pathogens such as *Staphylococcus aureus* which produces a small arsenal of inhibitors that function at the level of C3/C3b (44–49). Evaluating a potential synergy at the level of C1 between BBK32 and ElpB and/or ElpQ, as well as determining the contribution of ElpB and ElpQ in models of borreliosis, are important future questions. While the paralogous nature of the *elp* gene family presents challenges to commonly used genetic approaches in the field, a recently established CRISPR interference approach in *B. burgdorferi* may provide a future avenue to addressing such questions in murine models of Lyme disease (50).

Our previous work showed that ElpB and ElpQ blocks the cleavage of C2 and C4 by purified C1s (31). Surprisingly, however, we found that Elp proteins did not block cleavage of a small peptide substrate of C1s (31). Protease target selectivity aside, this feature of Elp inhibition is inherently different than BBK32’s active site-targeting mechanism (30). This prompted us to consider the possibility that Elp proteins were instead targeting exosites on C1s. Although ElpB and ElpQ also block cleavage of soluble C2 by C1s, detailed mapping of C2 exosites has not been described. In contrast, previous structural and biochemical studies have definitively mapped C4 exosites on C1s (6, 7, 9), and thus allowed us to test this hypothesis in detail.

Consistent with our hypothesis, we isolated an Elp binding site on activated C1s-CCP1-CCP2-SP, which contains both known C4 exosites (6, 7, 9) (i.e., CCPE and ABE) (**Fig. 3**). Importantly, ElpQ blocked C1s-CCP1-CCP2-SP-mediated C4 cleavage (**Fig. 4**) and C4 competed for ElpQ/C1s-CCP1-CCP2-SP_S632A_ interaction (**Fig. 5**). Furthermore, mutation of the four positively charged residues previously associated with the exosite on the SP domain (*i.e.* ABE) had a dramatic impact on the ability of ElpB and ElpQ to bind activated C1s-CCP1-CCP2-SP (**Fig. 6**). Collectively, our data strongly support a model in which ElpB and ElpQ target the ABE on C1s, and in this way Elp/C1s complex formation blocks C4 binding, and interferes with its cleavage by C1s. Given that the ABE is outside of the active site and forms completely only in activated C1s (**Fig. 6A**), this model is also consistent with the observed selectivity profile of Elp proteins for active C1s over zymogen C1s. While other bacterial protease inhibitors have been identified that target exosites, such as *E. coli* ecotin, this targeting is typically in tandem with active site blockade (51–53). In contrast to bacteria, a few naturally derived exosite-only inhibitors of the coagulation protease thrombin (Factor IIa) have been described from hematophagous organisms (reviewed in (54)). Our current model most resembles the exosite-only mechanisms used by triabin of the triatomine bug *Triatoma pallidipennis* (55) and C-terminal truncations of hirudin (*e.g.*, hirugen) from the medicinal leech *Hirudo medicinalis* (56), both of which interact with exosite I of thrombin and do not inhibit the cleavage of a small peptide.

Although we show here that targeting of C1s exosites is important for Elp activity, we also predict that the function of Elp proteins likely involves additional features. Of particular note is that each Elp protein has high affinity for purified C1r (31). Interestingly, however, and different from BBK32, binding of C1r by Elp proteins did not prevent cleavage of its natural substrate (*i.e.*, zymogen C1s), nor of small synthetic C1r peptidic substrates (31). This finding is consistent with the inhibitory mechanism proposed here, as C4 exosites are clearly absent in C1r. However, it also suggests that a common site on C1r and C1s exists that promotes affinity to each protease or that Elp proteins are capable of recognizing C1r at unique sites not found in C1s. Further studies will be needed, including elucidation of ElpB and ElpQ protein structures and/or complexes, to determine the role of C1r binding by this unusual class of protease inhibitor.

In summary, the unique inhibitory properties of ElpB and ElpQ described here provide new insights into microbial complement evasion by revealing an exosite-targeted, CP-selective, inhibitory mechanism present in Lyme disease spirochetes. This represents a novel form of complement evasion by a microbial pathogen and further study of this class of CP inhibitors may help guide future efforts to therapeutically target complement-mediated diseases.

## Experimental Procedures

### Plasmid construction

Elp constructs were PCR amplified with encoded 5’ BamHI and 3’ NotI restriction sites from the previously generated full-length ElpQ_19-343_ and ElpB_19-378_ pT7HMT plasmids using Q5 Master Mix (NEB) as previously described (31). DNA fragments for C1s constructs were *E. coli* codon optimized and synthetically produced by Integrated DNA Technologies gBlock Gene Fragment service with 5’ BamHI and 3’ NotI sites. All DNA fragments were then subjected to restriction enzyme digestion, ligated into pT7HMT (57), and transformed into *E. coli* DH5α as previously described (29, 30). Subsequent transformants were selected using LB plates containing 50 μg/mL kanamycin and sequence verified (Eurofins).

### Protein production and purification

GST-Elp and untagged Elp proteins were produced and purified as previously described (31). Briefly, following bacterial cell lysis, soluble fractions were subjected to nickel affinity purification (GoldBio) after being exchanged into native binding buffer (20 mM Tris (pH 8.0), 10 mM Imidazole, 500 mM NaCl) and eluted with (20 mM Tris (pH 8.0), 500 mM Imidazole, 500 mM NaCl). Eluted fractions were then exchanged into native binding buffer using a HiPrep Desalting 26/10 column on an ÄKTA pure FPLC (GE Healthcare). N-terminal tags were removed from Elp proteins using His-tagged tobacco etch virus (His-TEV) with the addition of 5 mM β-mercaptoethanol overnight at room temperature. Purification of the untagged Elp proteins proceeded by capturing His-TEV and uncleaved proteins on a HisTrap-FF 5mL column and collecting the flow through. Untagged Elp proteins were then further purified by size exclusion chromatography using a HiLoad 26/600 Superdex 75 PG gel filtration column (GE Healthcare) in HBS (10 mM HEPES (pH 7.3), 140 mM NaCl). Single peaks were concentrated, and samples were assessed for purity by SDS-PAGE analysis.

Human C1r enzyme, C1s enzyme, and C4 were purchased from Complement Technologies (Tyler, TX). Recombinant complement proteins corresponding to C1s were produced according to previously published methods (30, 58), with the following modifications: Tagged c-Myc-C1s-CCP1-CCP2-SP_S632A_ was not subjected to His-TEV cleavage and contains a c-Myc epitope as well as a 6X-His tag (57). C1s expresses as a zymogen and must be activated by treatment with C1r (59). When required, C1s SP domain containing proteins were incubated with 25 nM C1r enzyme overnight at 37° C in HBS-Ca^2+^ (10 mM HEPES (pH 7.3), 140 mM NaCl, 5 mM CaCl_2_). Activated C1s was then separated from C1r using an analytical size exclusion chromatography column Superdex 200 Increase 10/300 GL (GE Healthcare) in HBS. The peak containing activated C1s was then subjected to SDS-PAGE under reducing conditions and the presence of two chains was confirmed (6).

### ELISA-based complement inhibition assays

Pathway specific inhibition by Elp proteins was assessed by an ELISA-based approach as described elsewhere (28, 30, 31). Each pathway-specific initiator was immobilized on high-binding ELISA plates (Greiner Bio-One) with 3 μg ml^−1^ human IgM (CP; MP Biomedical), 20 μg ml^−1^ of mannan from *Saccharomyces cerevisiae* (LP; Sigma), or 25 μg ml^−1^ *Salmonella enteritidis* LPS (AP; Sigma) in coating buffer (100 mM Na_2_CO_3_/NaHCO_3_ (pH 9.6)). Reactions proceeded with 2% normal human serum (NHS) (CP; Innovative Research), 2% C1q-depleted serum (LP; Complement Technologies), or 20% NHS (AP; Innovative Research) and addition of 2 μM Elp proteins in either CP/LP buffer (10 mM HEPES (pH 7.3), 0.1% (w/v) gelatin, 140 mM NaCl, 2 mM CaCl_2_, 0.5 mM MgCl_2_) or AP buffer (10 mM HEPES (pH 7.3), 0.1% (w/v) gelatin, 140 mM NaCl, 5 mM MgCl_2_, 10 mM EGTA). Dose-dependent Elp inhibition of the classical pathway was determined using a 12-point, two-fold dilution (2200 – 1.1 nM) of Elp proteins. CP/LP activation was determined through C4b detection using mAb A211 (Quidel). AP activation was determined through C3c detection using mAb WM-1 (Sigma). All assays were performed in triplicate and normalized to positive (100%, no inhibitor) and negative (0%, no serum) controls. Inhibitory curves were fitted using non-linear regression to a normalized variable slope model in GraphPad v9 (Prism).

### Classical pathway hemolysis assay

Classical pathway specific inhibition of red blood cell lysis by Elp proteins were assessed by a hemolytic assay as described (28). In short, Elp truncations were added to sensitized sheep erythrocytes (Complement Technology) and 2% normal human serum (NHS; Innovative Research) in GHB^++^ (10 mM HEPES (pH 7.3), 140 mM NaCl, 0.1% gelatin (w/v), 0.15 mM CaCl_2_, and 0.5 mM MgCl_2_). Assays were performed in triplicate and normalized to positive (100%, no inhibitor) and negative (0%, no serum) controls. Inhibitory curves were fitted using non-linear regression to a normalized variable slope model in Graphpad v9 (Prism).

### Circular dichroism spectroscopy

Assessment of secondary structure for ElpQ_19-343_, ElpQ_181-343_, ElpB_19-378_, and ElpQ_182-378_ was performed using circular dichroism (CD) using previously described methods (43). Samples were diluted to 10 μM in 10 mM Na_3_PO_4_ and CD spectra were collected using a Chirascan V100 (Applied Photophysics, UK) with a square quartz cuvette with a path length of 0.05 cm across a wavelength range of 180-300 nm, at 120 nm min^−1^, using 1 nm step, 0.5 sec response, and 1 nm bandwidth. Spectra were background corrected against matching buffer using Prodata viewer (Applied Photophysics, UK).

### Surface plasmon resonance

SPR binding assays were carried out using general methods described previously (30, 31). Elp protein truncations were amine coupled to the CMD200 sensorchip (Xantec bioanalytics) using at 10 μg/ml in 10 mM sodium acetate pH 4.0 with final immobilization densities as follows: ElpQ_19-343_ (986 RU), ElpQ_181-343_ (311 RU), ElpQ_19-216_ (545 RU), ElpB_182-378_ (575 RU), ElpB_111-378_ (554 RU), ElpB_288-378_ (473 RU). All assays were performed in a running buffer of HBS-T-Ca^2+^ (10 mM HEPES (pH 7.3), 140 mM NaCl, 0.005% (v/v) Tween 20, 5 mM CaCl_2_) with a flowrate of 30 μl min^−1^. Analytes were exchanged into matching running buffer prior to experimentation and after each analyte injection surfaces were regenerated to baseline using three 60 sec injections of 2M NaCl. Elp interaction with recombinant C1s domain truncations was evaluated using single injections of each C1s truncation at 500 nM with an association time of 120 sec and a dissociation time of 120 sec or by multi-cycle experiments performed with C1s enzyme (Complement Technology) as well as, recombinant wild-type C1s-CCP1-CCP2-SP or exosite mutants using the injection series (200, 100, 50, 25, 12.5, 6.3, 3.1, 1.6, 0.8, and 0 nM) over an association time of 120 sec followed by a 180 sec dissociation. All injection series were performed in triplicate. Kinetic (*K*_D, kin_) and steady-state (*K*_D, ss_) fits of the sensorgrams were determined using a 1:1 binding model (Langmuir) using Biacore T200 Evaluation Software 3.1 (GE Healthcare).

### SDS-PAGE-based C4 cleavage assay

C1s cleavage of C4 in the presence of ElpQ was assessed using SDS-PAGE analysis as previously described (31). Briefly, a 10 μL reaction in HBS-Ca^2+^ (10 mM HEPES (pH 7.3), 140 mM NaCl, 5 mM CaCl_2_) of recombinant activated C1s-CCP1-CCP2-SP at 6.25 nM and two-fold dilutions of ElpQ_182-343_ starting at 25,000 nM was added to 1.25μL of C4 (1 mg/mL) (Complement Technologies). The reaction was incubated at 37° C for 1 hr and was stopped by addition of 5 μL Laemmli buffer and subsequently boiled for 5 min. Coomassie staining was performed and experimentation was performed in triplicate. Densitometric analysis of the gels proceeded using Image Lab™ (Bio-Rad) with lanes and bands manually selected. Intensity of bands corresponding to C4α’ (~85 kDa) were in lane normalized to C4β (~75 kDa) and plotted with 100% cleavage defined as the positive control lane C1s + C4 and 0% as C4 only. Quantification of IC_50_ was analyzed using a four-parameter nonparametric response in GraphPad v9.3 (Prism).

### C1s Enzyme assay

Active site inhibition of C1s by ElpQ was measured using a chromogenic substrate cleavage assay as previously described using the following modifications (31). ElpQ_181-343_ or Futhan (Sigma), a broad specificity complement protease inhibitor, at a concentration of 25 μM was added to 100 μM Z-L-Lys thiobenzyl (Sigma) and 100 μM DTNB (TCI) in HBS-Ca^2+^ (10 mM HEPES (pH 7.3), 140 mM NaCl, 5 mM CaCl_2_). Just prior to measurement, 6.25 nM C1s-CCP1-CCP2-SP was added for a final 80 μl reaction volume. Reactions were carried out at 37° C for 1 hr and read at 412 nm using a Versamax plate reader (Molecular Devices) in triplicate. Data were normalized by including C1s with substrate as a 100% cleavage control, or peptide and DTNB as 0%.

### Alpha competition experiments

Competitive binding disrupting the ElpQ-C1s interaction was assessed using an <u>a</u>mplified <u>l</u>uminescence <u>p</u>roximity <u>h</u>omogenous <u>a</u>ssay (Alpha). First, optimal concentrations of the donor protein (GST-ElpQ) and acceptor protein (c-Myc-C1s-CCP1-CCP2-SP_S632A_) was determined with an 8 x 8 cross-titration experiment (**Fig S4**). A two-fold concentration series (300 – 4.7 nM) of c-Myc-C1s-CCP1-CCP2-SP_S632A_ and (100 – 1.6 nM) GST-ElpQ were diluted in HBS-T Ca^2+^ (10 mM HEPES (pH 7.3), 140 mM NaCl, 0.005% (v/v) Tween 20, 5 mM CaCl_2_). Addition of 20 μg mL^−1^ Glutathione-Donor bead (PerkinElmer) and lastly 20 μg mL^−1^ Anti-c-Myc AlphaLISA^®^ Acceptor bead (PerkinElmer) completed the total reaction volume of 25 μL. Steady-state binding signal was obtained after incubation of reactions for 1 hr at 25° C using a EnSight Multimode plate reader (PerkinElmer) with a 30 ms excitation at 680 nm and emission measured at 615 nm after 140 ms. Subsequent single experimentation with 500 nM C1s-CCP1-CCP2-SP_S632A_ and 50 nM GST-ElpQ using the same bead concentrations yielded excellent signal to noise and follows the Cheng-Prusoff equation where the IC_50_ approximates the *K*_D_ when there is a ten-fold difference in concentration of the target (ElpQ) versus the tracer (C1s) (60).

Competition assays were performed using 12-point two-fold dilution with top concentrations of 6,100 nM ElpQ_19-343_, 1,000 nM C4 (Complement Technologies), 2,000 nM ElpQ_181-343_, or 2,000 nM ElpB_182-378_ with the c-Myc acceptor bead added last. Alpha signal (counts) were normalized to the absence of competitor (100%) and absence of c-Myc-C1s-CCP1-CCP2-SP S632A (0%) controls. All experiments were performed in triplicate and fitted using non-linear regression to a normalized variable slope model in Graphpad v9 (Prism).

## Data Availability

All data are contained within the manuscript.

## Supporting Information

This article contains supporting information.

## Author Contributions

R.J.G., J.M.L. and B.L.G. designed research; R.J.G. and S.T. conducted research; R.J.G., S.T., J.M.L. and B.L.G analyzed data; R.J.G. and B.L.G. wrote the paper.

## Funding and additional information

Support was provided by Public Health Service Grant R01-AI146930 from the National Institute of Allergy and Infectious Diseases (B.L.G). The content is solely the responsibility of the authors and does not necessarily represent the official views of the National Institutes of Health

## Abbreviations

CP: classical pathway
LP: lectin pathway
AP: alternative pathway
MBL: mannan-binding lectin
MASP: mannan-binding lectin associated protease
CUB: C1r/C1s, Uegf, Bmp1
EGF: epidermal growth factor-like
CCP: complement control protein
SP: serine protease
ABE: anion-binding exosite
CCPE: complement control protein domain exosite
RCA: regulator of complement activity
SPR: surface plasmon resonance
*K*_D_: equilibrium dissociation constant
SS: steady state
Alpha: amplified luminescence proximity homogenous assay
NHS: normal human serum
CD: circular dichroism

## References

1. Merle, N. S., Church, S. E., Fremeaux-Bacchi, V., and Roumenina, L. T. (2015) Complement System Part I – Molecular Mechanisms of Activation and Regulation. Front Immunol. 6, 262

2. Merle, N. S., Noe, R., Halbwachs-Mecarelli, L., Fremeaux-Bacchi, V., and Roumenina, L. T. (2015) Complement System Part II: Role in Immunity. Front Immunol. 6, 257

3. Walport, M. J. (2001) Complement. First of two parts. The New England Journal of Medicine. 344, 1058–1066

4. Walport, M. J. (2001) Complement. Second of two parts. The New England Journal of Medicine. 344, 1140–1144

5. Schechter, I., and Berger, A. (1967) On the size of the active site in proteases. I. Papain. Biochemical and Biophysical Research Communications. 27, 157–162

6. Duncan, R. C., Mohlin, F., Taleski, D., Coetzer, T. H., Huntington, J. A., Payne, R. J., Blom, A. M., Pike, R. N., and Wijeyewickrema, L. C. (2012) Identification of a catalytic exosite for complement component C4 on the serine protease domain of C1s. J Immunol. 189, 2365–2373

7. Perry, A. J., Wijeyewickrema, L. C., Wilmann, P. G., Gunzburg, M. J., D’Andrea, L., Irving, J. A., Pang, S. S., Duncan, R. C., Wilce, J. A., Whisstock, J. C., and Pike, R. N. (2013) A molecular switch governs the interaction between the human complement protease C1s and its substrate, complement C4. The Journal of Biological Chemistry. 288, 15821–15829

8. Gaboriaud, C., Rossi, V., Bally, I., Arlaud, G. J., and Fontecilla-Camps, J. C. (2000) Crystal structure of the catalytic domain of human complement c1s: a serine protease with a handle. EMBO J. 19, 1755–1765

9. Kidmose, R. T., Laursen, N. S., Dobó, J., Kjaer, T. R., Sirotkina, S., Yatime, L., Sottrup-Jensen, L., Thiel, S., Gál, P., and Andersen, G. R. (2012) Structural basis for activation of the complement system by component C4 cleavage. Proc Natl Acad Sci U S A. 109, 15425–15430

10. Bock, P. E., Panizzi, P., and Verhamme, I. M. A. (2007) Exosites in the substrate specificity of blood coagulation reactions. J Thromb Haemost. 5 Suppl 1, 81–94

11. Schmidt, C. Q., Lambris, J. D., and Ricklin, D. (2016) Protection of host cells by complement regulators. Immunological Reviews. 274, 152–171

12. Ricklin, D., and Lambris, J. D. (2013) Complement in Immune and Inflammatory Disorders: Therapeutic Interventions. The Journal of Immunology. 190, 3839–3847

13. Lambris, J. D., Ricklin, D., and Geisbrecht, B. v. (2008) Complement evasion by human pathogens. Nature Reviews Microbiology. 6, 132–142

14. Garcia, B. L., Zwarthoff, S. A., Rooijakkers, S. H. M., and Geisbrecht, B. v (2016) Novel Evasion Mechanisms of the Classical Complement Pathway. Journal of Immunology. 197, 2051–2060

15. Kugeler, K. J., Schwartz, A. M., Delorey, M. J., Mead, P. S., and Hinckley, A. F. (2021) Estimating the Frequency of Lyme Disease Diagnoses, United States, 2010-2018. Emerg Infect Dis. 27, 616–619

16. Skare, J. T., and Garcia, B. L. (2020) Complement Evasion by Lyme Disease Spirochetes. Trends in Microbiology. 28, 889–899

17. Coburn, J., Garcia, B., Hu, L. T., Jewett, M. W., Kraiczy, P., Norris, S. J., and Skare, J. (2021) Lyme Disease Pathogenesis. Curr Issues Mol Biol. 42, 473–518

18. Kraiczy, P., and Stevenson, B. (2013) Complement regulator-acquiring surface proteins of Borrelia burgdorferi: Structure, function and regulation of gene expression. Ticks and Tick-Borne Diseases. 4, 26–34

19. Lin, Y. P., Frye, A. M., Nowak, T. A., and Kraiczy, P. (2020) New Insights Into CRASP-Mediated Complement Evasion in the Lyme Disease Enzootic Cycle. Front Cell Infect Microbiol. 10.3389/FCIMB.2020.00001

20. Kraiczy, P., Skerka, C., Kirschfink, M., Zipfel, P. F., and Brade, V. (2001) Mechanism of complement resistance of pathogenic *Borrelia burgdorferi* isolates. International Immunopharmacology. 1, 393–401

21. Hartmann, K., Corvey, C., Skerka, C., Kirschfink, M., Karas, M., Brade, V., Miller, J. C., Stevenson, B., Wallich, R., Zipfel, P. F., and Kraiczy, P. (2006) Functional characterization of BbCRASP-2, a distinct outer membrane protein of *Borrelia burgdorferi* that binds host complement regulators factor H and FHL-1. Molecular Microbiology. 61, 1220–1236

22. McDowell, J. v, Hovis, K. M., Zhang, H., Tran, E., Lankford, J., and Marconi, R. T. (2006) Evidence that the BBA68 protein (BbCRASP-1) of the Lyme disease spirochetes does not contribute to factor H-mediated immune evasion in humans and other animals. Infection and Immunity. 74, 3030–3034

23. McDowell, J. v., Wolfgang, J., Tran, E., Metts, M. S., Hamilton, D., and Marconi, R. T. (2003) Comprehensive analysis of the factor H binding capabilities of *Borrelia* species associated with lyme disease: Delineation of two distinct classes of factor H binding proteins. Infection and Immunity. 71, 3597–3602

24. Bykowski, T., Woodman, M. E., Cooley, A. E., Brissette, C. A., Wallich, R., Brade, V., Kraiczy, P., and Stevenson, B. (2008) *Borrelia burgdorferi* complement regulator-acquiring surface proteins (BbCRASPs): Expression patterns during the mammal-tick infection cycle. International Journal of Medical Microbiology. 298, 249–256

25. Kenedy, M. R., Vuppala, S. R., Siegel, C., Kraiczy, P., and Akins, D. R. (2009) CspA-mediated binding of human factor H inhibits complement deposition and confers serum resistance in *Borrelia burgdorferi*. Infection and Immunity. 77, 2773–2782

26. Rogers, E. A., Abdunnur, S. v., McDowell, J. v., and Marconi, R. T. (2009) Comparative analysis of the properties and ligand binding characteristics of CspZ, a factor H binding protein, derived from *Borrelia burgdorferi* isolates of human origin. Infection and Immunity. 77, 4396–4405

27. Hallström, T., Siegel, C., Mörgelin, M., Kraiczy, P., Skerka, C., and Zipfel, P. F. (2013) CspA from *Borrelia burgdorferi* inhibits the terminal complement pathway. mBio. 4:e00481, 13

28. Garcia, B. L., Zhi, H., Wager, B., Höök, M., and Skare, J. T. (2016) *Borrelia burgdorferi* BBK32 Inhibits the Classical Pathway by Blocking Activation of the C1 Complement Complex. PLoS Pathogens. 12, e1005404

29. Xie, J., Zhi, H., Garrigues, R. J., Keightley, A., Garcia, B. L., and Skare, J. T. (2019) Structural determination of the complement inhibitory domain of *Borrelia burgdorferi* BBK32 provides insight into classical pathway complement evasion by Lyme disease spirochetes. PLoS Pathog. 15, e1007659

30. Garrigues, R. J., Powell-Pierce, A. D., Hammel, M., Skare, J. T., and Garcia, B. L. (2021) A Structural Basis for Inhibition of the Complement Initiator Protease C1r by Lyme Disease Spirochetes. J Immunol. 207, 2856–2867

31. Pereira, M. J., Wager, B., Garrigues, R. J., Gerlach, E., Quinn, J. D., Dowdell, A. S., Osburne, M. S., Zückert, W. R., Kraiczy, P., Garcia, B. L., and Leong, J. M. (2022) Lipoproteome screening of the Lyme disease agent identifies inhibitors of antibody-mediated complement killing. Proc Natl Acad Sci U S A. 119, e2117770119

32. Dowdell, A. S., Murphy, M. D., Azodi, C., Swanson, S. K., Florens, L., Chen, S., and Zückert, W. R. (2017) Comprehensive spatial analysis of the *Borrelia burgdorferi* lipoproteome reveals a compartmentalization bias toward the bacterial surface. Journal of Bacteriology. 199, E00658

33. Stevenson, B., Tilly, K., and Rosa, P. A. (1996) A family of genes located on four separate 32-kilobase circular plasmids in *Borrelia burgdorferi* B31. Journal of Bacteriology. 178, 3508–3516

34. Akins, D. R., Caimano, M. J., Yang, X., Cerna, F., Norgard, M. v., and Radolf, J. D. (1999) Molecular and evolutionary analysis of *Borrelia burgdorferi* 297 circular plasmid-encoded lipoproteins with OspE– and OspF-like leader peptides. Infection and Immunity. 10.1128/iai.67.3.1526-1532.1999

35. Marconi, R. T., Sung, S. Y., Hughes, C. A. N., and Carlyon, J. A. (1996) Molecular and evolutionary analyses of a variable series of genes in *Borrelia burgdorferi* that are related to ospE and ospF, constitute a gene family, and share a common upstream homology box. Journal of Bacteriology. 10.1128/jb.178.19.5615-5626.1996

36. Brissette, C. A., Cooley, A. E., Burns, L. H., Riley, S. P., Verma, A., Woodman, M. E., Bykowski, T., and Stevenson, B. (2008) Lyme borreliosis spirochete Erp proteins, their known host ligands, and potential roles in mammalian infection. Int J Med Microbiol. 298 Suppl, 257–67

37. Stevenson, B., Krusenstjerna, A. C., Castro-Padovani, T. N., Savage, C. R., Jutras, B. L., and Saylor, T. C. (2022) The Consistent Tick-Vertebrate Infectious Cycle of the Lyme Disease Spirochete Enables *Borrelia burgdorferi* To Control Protein Expression by Monitoring Its Physiological Status. J Bacteriol. 10.1128/jb.00606-21

38. Brissette, C. A., Verma, A., Bowman, A., Cooley, A. E., and Stevenson, B. (2009) The *Borrelia burgdorferi* outer-surface protein ErpX binds mammalian laminin. Microbiology (Reading). 155, 863–872

39. Stevenson, B., Bono, J. L., Schwan, T. G., and Rosa, P. (1998) *Borrelia burgdorferi* Erp proteins are immunogenic in mammals infected by tick bite, and their synthesis is inducible in cultured bacteria. Infection and Immunity. 10.1128/iai.66.6.2648-2654.1998

40. Bielefeld-Sevigny, M. (2009) AlphaLISA immunoassay platform– the “no-wash” high-throughput alternative to ELISA. Assay Drug Dev Technol. 7, 90–92

41. Zückert, W. R. (2014) Secretion of Bacterial Lipoproteins: Through the Cytoplasmic Membrane, the Periplasm and Beyond. Biochimica et Biophysica Acta (BBA) – Molecular Cell Research. 1843, 1509–1516

42. Zückert, W. R. (2019) Protein Secretion in Spirochetes. Microbiol Spectr. 10.1128/MICROBIOLSPEC.PSIB-0026-2019

43. Booth, C. E., Powell-Pierce, A. D., Skare, J. T., and Garcia, B. L. (2022) Borrelia miyamotoi FbpA and FbpB Are Immunomodulatory Outer Surface Lipoproteins With Distinct Structures and Functions. Front Immunol. 10.3389/FIMMU.2022.886733

44. Clark, E. A., Crennell, S., Upadhyay, A., Zozulya, A. v., Mackay, J. D., Svergun, D. I., Bagby, S., and van den Elsen, J. M. H. (2011) A structural basis for Staphylococcal complement subversion: X-ray structure of the complement-binding domain of Staphylococcus aureus protein Sbi in complex with ligand C3d. Mol Immunol. 48, 452–462

45. Hammel, M., Sfyroera, G., Pyrpassopoulos, S., Ricklin, D., Ramyar, K. X., Pop, M., Jin, Z., Lambris, J. D., and Geisbrecht, B. v. (2007) Characterization of Ehp, a secreted complement inhibitory protein from *Staphylococcus aureus*. J Biol Chem. 282, 30051–30061

46. Hammel, M., Sfyroera, G., Ricklin, D., Magotti, P., Lambris, J. D., and Geisbrecht, B. v (2007) A structural basis for complement inhibition by *Staphylococcus aureus*. Nature Immunology. 8, 430–437

47. Ricklin, D., Ricklin-Lichtsteiner, S. K., Markiewski, M. M., Geisbrecht, B. v., and Lambris, J. D. (2008) Cutting edge: members of the *Staphylococcus aureus* extracellular fibrinogen-binding protein family inhibit the interaction of C3d with complement receptor 2. J Immunol. 181, 7463–7467

48. Amdahl, H., Jongerius, I., Meri, T., Pasanen, T., Hyvärinen, S., Haapasalo, K., van Strijp, J. A., Rooijakkers, S. H., and Jokiranta, T. S. (2013) Staphylococcal Ecb protein and host complement regulator factor H enhance functions of each other in bacterial immune evasion. J Immunol. 191, 1775–1784

49. Rooijakkers, S. H. M., Wu, J., Ruyken, M., van Domselaar, R., Planken, K. L., Tzekou, A., Ricklin, D., Lambris, J. D., Janssen, B. J. C., van Strijp, J. A. G., and Gros, P. (2009) Structural and functional implications of the alternative complement pathway C3 convertase stabilized by a staphylococcal inhibitor. Nature Immunology. 10, 721–727

50. Takacs, C. N., Scott, M., Chang, Y., Kloos, Z. A., Irnov, I., Rosa, P. A., Liu, J., and Jacobs-Wagner, C. (2020) A CRISPR interference platform for selective downregulation of gene expression in *Borrelia burgdorferi*. Appl Environ Microbiol. 87, e02519–20

51. Gaboriaud, C., Gupta, R. K., Martin, L., Lacroix, M., Serre, L., Teillet, F., Arlaud, G. J., Rossi, V., and Thielens, N. M. (2013) The serine protease domain of MASP-3: enzymatic properties and crystal structure in complex with ecotin. PloS One. 8, e67962

52. Nagy, Z. A., Héja, D., Bencze, D., Kiss, B., Boros, E., Szakács, D., Fodor, K., Wilmanns, M., Kocsis, A., Dobó, J., Gál, P., Harmat, V., and Pál, G. (2022) Synergy of protease-binding sites within the ecotin homodimer is crucial for inhibition of MASP enzymes and for blocking lectin pathway activation. J Biol Chem. 10.1016/J.JBC.2022.101985

53. Farady, C. J., and Craik, C. S. (2010) Mechanisms of macromolecular protease inhibitors. Chembiochem. 11, 2341–2346

54. Troisi, R., Balasco, N., Autiero, I., Vitagliano, L., and Sica, F. (2021) Exosite Binding in Thrombin: A Global Structural/Dynamic Overview of Complexes with Aptamers and Other Ligands. International Journal of Molecular Sciences 2021, *Vol. 22, Page 10803*. 22, 10803

55. Fuentes-Prior, P., Noeske-Jungblut, C., Donner, P., Schleuning, W. D., Huber, R., and Bode, W. (1997) Structure of the thrombin complex with triabin, a lipocalin-like exosite-binding inhibitor derived from a triatomine bug. Proc Natl Acad Sci U S A. 94, 11845–11850

56. Skrzypczak-Jankun, E., Carperos, V. E., Ravichandran, K. G., Tulinsky, A., Westbrook, M., and Maraganore, J. M. (1991) Structure of the hirugen and hirulog 1 complexes of α-thrombin. Journal of Molecular Biology. 221, 1379–1393

57. Geisbrecht, B. v, Bouyain, S., and Pop, M. (2006) An optimized system for expression and purification of secreted bacterial proteins. Protein Expression and Purification. 46, 23–32

58. Rushing, B. R., Rohlik, D. L., Roy, S., Skaff, D. A., and Garcia, B. L. (2020) Targeting the Initiator Protease of the Classical Pathway of Complement Using Fragment-Based Drug Discovery. Molecules. 10.3390/molecules25174016

59. Rossi, V., Bally, I., Thielens, N. M., Esser, A. F., and Arlaud, G. J. (1998) Baculovirus-mediated Expression of Truncated Modular Fragments from the Catalytic Region of Human Complement Serine Protease C1s: EVIDENCE FOR THE INVOLVEMENT OF BOTH COMPLEMENT CONTROL PROTEIN MODULES IN THE RECOGNITION OF THE C4 PROTEIN SUBSTRATE. Journal of Biological Chemistry. 273, 1232–1239

60. Yung-Chi, C., and Prusoff, W. H. (1973) Relationship between the inhibition constant (K1) and the concentration of inhibitor which causes 50 per cent inhibition (I50) of an enzymatic reaction. Biochem Pharmacol. 22, 3099–3108

61. Madeira, F., Pearce, M., Tivey, A. R. N., Basutkar, P., Lee, J., Edbali, O., Madhusoodanan, N., Kolesnikov, A., and Lopez, R. (2022) Search and sequence analysis tools services from EMBL-EBI in 2022. Nucleic Acids Res. 10.1093/NAR/GKAC240

62. Robert, X., and Gouet, P. (2014) Deciphering key features in protein structures with the new ENDscript server. Nucleic Acids Res. 10.1093/NAR/GKU316

63. Jumper, J., Evans, R., Pritzel, A., Green, T., Figurnov, M., Ronneberger, O., Tunyasuvunakool, K., Bates, R., Žídek, A., Potapenko, A., Bridgland, A., Meyer, C., Kohl, S. A. A., Ballard, A. J., Cowie, A., Romera-Paredes, B., Nikolov, S., Jain, R., Adler, J., Back, T., Petersen, S., Reiman, D., Clancy, E., Zielinski, M., Steinegger, M., Pacholska, M., Berghammer, T., Bodenstein, S., Silver, D., Vinyals, O., Senior, A. W., Kavukcuoglu, K., Kohli, P., and Hassabis, D. (2021) Highly accurate protein structure prediction with AlphaFold. Nature. 596, 583–589

