## Supplemental Information for "Outer surface lipoproteins from the Lyme disease spirochete exploit the molecular switch mechanism of the complement protease C1s"

Running Title: *B. burgdorferi* C1s inhibitors

Ryan J. Garrigues[1], Sheila Thomas[1], John M. Leong[2], and Brandon L. Garcia[1]

[1]Department of Microbiology and Immunology, Brody School of Medicine, East Carolina University, Greenville, North Carolina, United States of America

[2]Department of Molecular Biology and Microbiology, Tufts School of Medicine, Tufts University, Boston, Massachusetts, USA

**This file contains Figures S1 – S5**

A

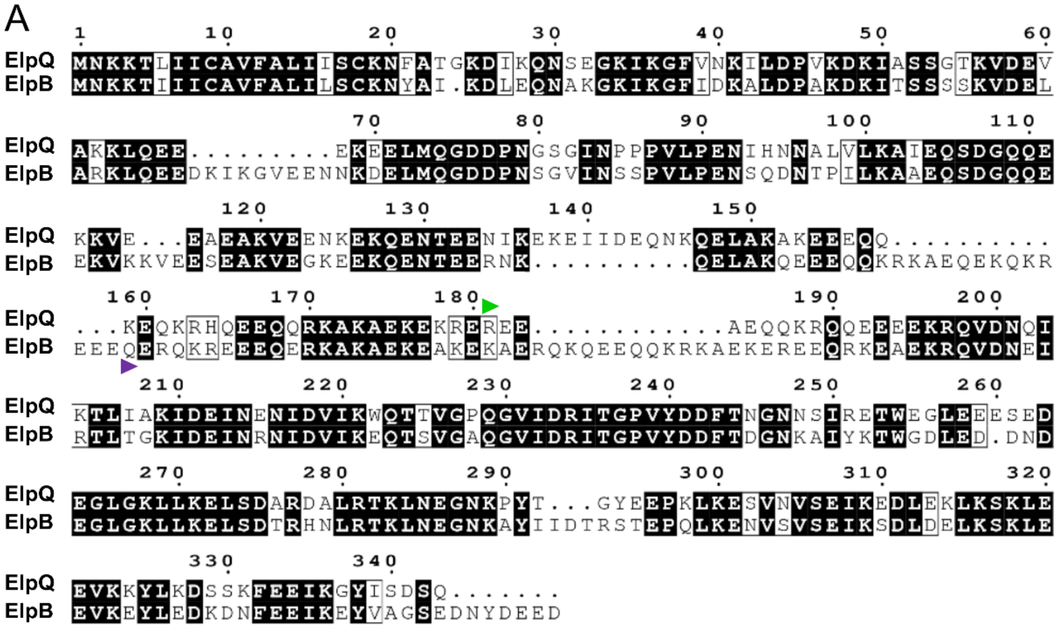


B

| | ElpB | ElpB$_{182-378}$ |
|---|---|---|
| ElpQ | 58 (*73*) | |
| ElpQ$_{181-343}$ | | 55 (*66*) |

**% Identity (*% Similarity*)**

**Figure S1.** Sequence alignment and homology of ElpB and ElpQ. **A)** Amino acid sequences of ElpB (UNIPROT #: H7C7R2) and ElpQ (UNIPROT #: Q9S035) from *B. burgdorferi* strain B31 were aligned using EMBOSS Needle and graphically modified using ESPript 3.0 (61, 62). Purple (ElpB) or green (ElpQ) arrows denote the N-terminal most residue in the minimal inhibitory constructs identified in **Fig. 1** (*i.e.* ElpB$_{182-378}$ and ElpQ$_{181-343}$). **B)** Percent identity and similarity were calculated for the full-length and minimal inhibitory domains.

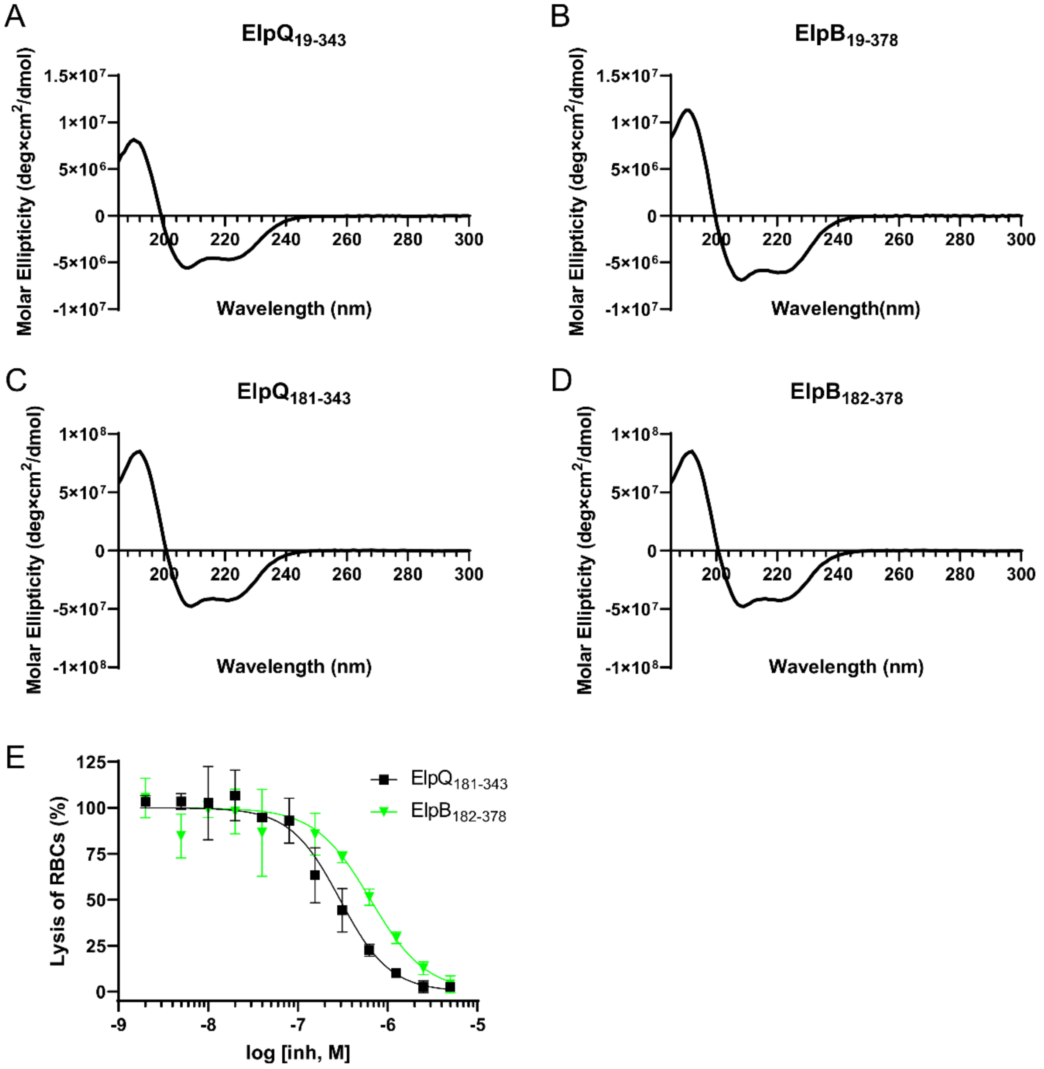


**Figure S2.** Circular dichroism spectroscopy. **A-D)** CD spectra indicate that ElpQ and ElpB and their associated minimal inhibitory domains, $ElpQ_{181-343}$ and $ElpB_{182-378}$, are folded and have predominantly alpha helical secondary structure. **E)** $ElpQ_{181-343}$ and $ElpB_{182-378}$ dose-dependently inhibited complement-mediated lysis of opsonized sheep red blood cells. Experiments were performed in duplicate. Non-linear regression analysis was performed to calculate $IC_{50}$ values: $ElpQ_{181-343}$ $IC_{50}$ = 300 nM, 95% confidence interval = 270 to 320 nM; $ElpB_{182-378}$ $IC_{50}$ = 650 nM, 95% confidence interval = 560 to 740 nM.

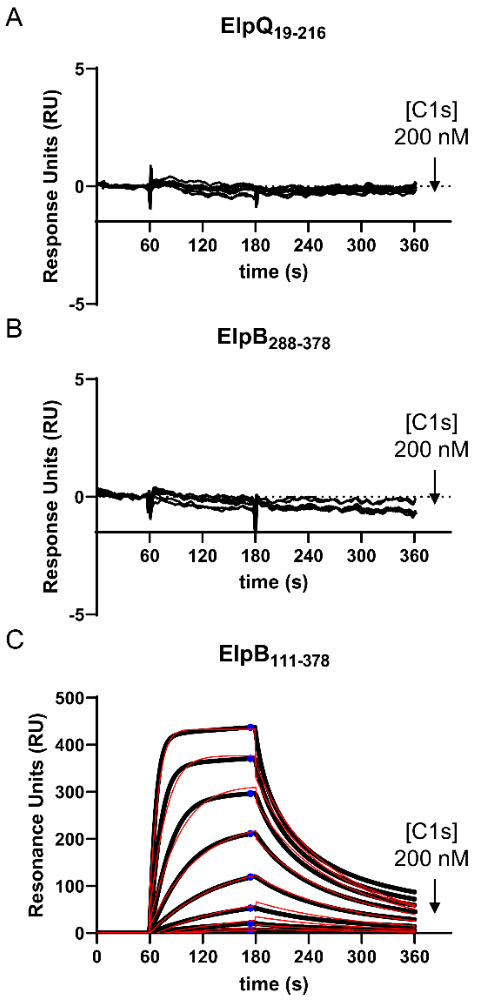


**Figure S3.** SPR binding assays using full-length C1s. **A-B)** Two constructs that lacked inhibitory activity in the CP ELISA assays (**Fig. 1**) were selected for C1s interaction analysis by SPR. Experiments were carried out in an identical manner as to those conducted for the minimal inhibitory constructs (**Fig. 2**). $ElpQ_{19-216}$ and $ElpB_{288-378}$ did not produce a detectable binding signal at C1s protein concentrations up to 200 nM. **C)** A longer C-terminal construct that exhibited full inhibitory activity in the CP ELISA was selected for C1s interaction analysis, $ElpB_{111-378}$. Kinetic fits are shown as red traces and steady-state responses are marked by blue circles. Representative injection series are shown. Each injection series was performed in triplicate.

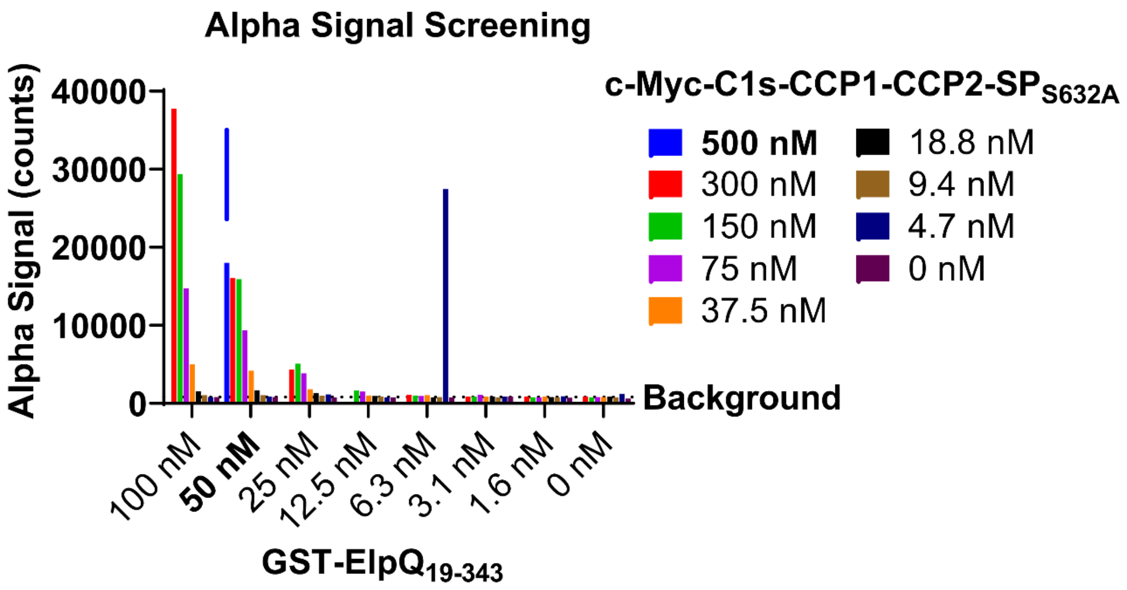


**Figure S4.** Alpha signal screening. Initial optimization of Alpha signal was performed using an 8 x 8 grid screen using 20 µg $mL^{-1}$ Anti-c-Myc-Acceptor bead, 20 µg $mL^{-1}$ glutathione-conjugated Donor bead, and twofold dilutions of c-Myc-C1s-CCP1-CCP2-SP and GST-ElpQ at top concentrations of 300 nM and 100 nM, respectively. A subsequent optimization reaction containing 500 nM c-Myc-C1s-CCP1-CCP2-SP and 50 nM GST-ElpQ (blue arrow) was selected for later competition-based experiments (see *Experimental Procedures*) (**Figs. 3D and 5**).

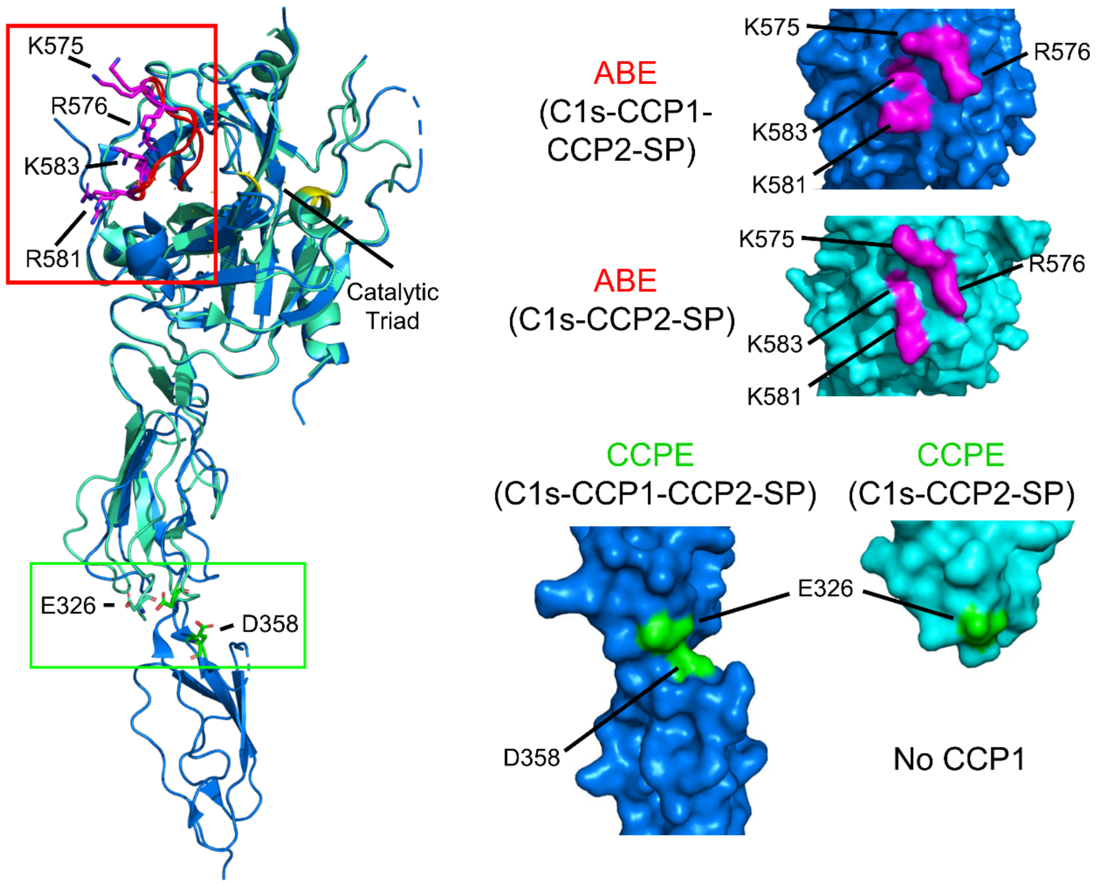


**Figure S5.** A structure of an unbound form of activated C1s-CCP1-CCP2-SP has not been reported, however this construct was used in the co-crystallization of gigastasin (PDB: 5UBM). Thus, the 5UBM structure is presented in **Fig. 6A** as a model for activated CCP1-CCP2-SP so that both ABE (red loop, magenta sticks) and CCPE (green sticks) exosites could be highlighted. The position and surface topology of residues involved in C4 binding in the apo form of activated C1s-CCP2-SP (cyan, PDB:1ELV) is similar to that of the Gigastasin-inhibited C1s-CCP1-CCP2-SP structure (blue, PDB: 5UBM).
